# Novel Master Regulators of Microglial Phagocytosis and Repurposed FDA-approved Drug for Treatment of Alzheimer Disease

**DOI:** 10.1101/2022.10.19.512953

**Authors:** Kuixi Zhu, Qianying He, Sheng-Feng Tsai, Dinusha Maheepala Mudalige, Marc Y.R. Henrion, Syed S.A. Zaidi, Lau Branden, Andrew Tang, Mika P. Cadiz, Rachel Hodos-Nkhereanye, Sara Moein, Melissa L. Alamprese, David A. Bennett, Philip L. De Jager, John D. Frye, Nilu□fer Ertekin-Taner, Yu-Min Kuo, Patrick T. Ronaldson, Rui Chang

**Affiliations:** The Center for Innovation in Brain Sciences, University of Arizona, Tucson, AZ, USA; Department of Biosystems Engineering, University of Arizona, Tucson, AZ, USA; Department of Cell Biology and Anatomy, College of Medicine, National Cheng Kung University, Tainan, Taiwan; Liverpool School of Tropical Medicine, Pembroke Place, Liverpool, Pembroke Place, L3 5QA, UK; Malawi - Liverpool - Wellcome Trust Clinical Research Programme, PO Box 30096, Blantyre, Malawi; Arizona Research Labs, Genetics Core, University of Arizona, Tucson, AZ, USA; Neuroscience Graduate Interdisciplinary Program, University of Arizona, Tucson, AZ, USA; Department of Neuroscience, Mayo Clinic, Scottsdale, AZ, 85259, USA; Neuroscience Graduate Program, Mayo Clinic Graduate School of Biomedical Sciences, Scottsdale, AZ, 85259, USA; Icahn School of Medicine at Mount Sinai, New York, NY, USA; BenevolentAI, Brooklyn, NY, USA; Department of Pathology and Laboratory Medicine, Weill Cornell Medicine, New York, NY, USA; Rush Alzheimer’s Disease Center, Rush University Medical Center, Chicago, IL, USA; Center for Translational & Computational Neuroimmunology, Department of Neurology, Columbia University Medical Center, New York, NY, USA; Neurodegeneration Program, New York Genome Center, NY, USA; Department of Neuroscience, Mayo Clinic Florida, Jacksonville, FL, USA; Department of Neurology, Mayo Clinic Florida, Jacksonville, FL, USA; Department of Pharmacology, College of Medicine, University of Arizona, Tucson, AZ, USA; Department of Neurology, University of Arizona, Tucson, AZ, USA; INTelico Therapeutics LLC, Tucson, AZ, USA; PATH Biotech LLC, Tucson, AZ, USA

**Author notes:** Authors equally contributed to the manuscript (Kuixi Zhu, Qianying He, Sheng-Feng Tsai, Dinusha Maheepala Mudalige).

## Abstract

Microglia, the innate immune cells of the brain, are essential determinants of late-onset Alzheimer’s Disease (LOAD) neuropathology. Here, we developed an integrative computational systems biology approach to construct causal network models of genetic regulatory programs for microglia in Alzheimer’s Disease (AD). This model enabled us to identify novel key driver (KDs) genes for microglial functions that can be targeted for AD pharmacotherapy. We prioritized *FCER1G, HCK, LAPTM5, ITGB2, SLC1A2, PAPLN, GSAP, NTRK2*, and *CIRBP* as KDs of microglial phagocytosis promoting neuroprotection and/or neural repair. *In vitro*, shRNA knockdown of each KD significantly reduced microglial phagocytosis. We repurposed riluzole, an FDA-approved ALS drug that upregulates *SLC1A2* activity, and discovered that it stimulated phagocytosis of Aβ1-42 in human primary microglia and decreased hippocampal amyloid plaque burden/phosphorylated tau levels in the brain of aged 3xTg-AD mice. Taken together, these data emphasize the utlility of our integrative approach for repurposing drugs for AD therapy.

## Introduction

Late-onset Alzheimer’s disease (LOAD) is a complex neurodegenerative disease that is characterized by neuropathology consisting of amyloid plaques, neurofibrillary tangles, and clinical dementia. Genome-wide association studies (GWAS) have implicated immune cell-specific genes associated with AD risk and point to microglia as a causal cell type[1-3]. Microglial cells are resident innate immune cells of the central nervous system (CNS) [4] and play a critical role in abolishing apoptotic cells, Aβ deposits and synapse removal by phagocytosis [5]. It has previously been discovered that microglia are colocalized with amyloid plaques in the brain [6]. Given the recently failed clinical trials directly targeting anti-amyloid plaques [7], it becomes essential to understand molecular mechanisms of microglia in formation and clearance of amyloid plaques. A previous study using co-expression network analysis of post-mortem brain tissue from patients with a positive diagnosis of AD showed a microglial module dominated by genes implicated in phagocytosis [8]. Indeed, the neuroprotective function of microglia-mediated phagocytosis engulfing dead/dying neurons and neuronal debris under normal physiological conditions have been appreciated for many years because it is a vital process involved in mitigation of neuroinflammation [9]. Therefore, it is critical to improve our understanding of these discrete molecular mechanisms of microglial activation and phagocytosis, which will facilitate development of novel therapeutics. This objective can be accomplished by identification of potential master regulators (key drivers, KDs) that modulate microglial functions, particularly those involved in phagocytosis and Aβ clearance in the setting of AD.

## Results

### Integrative Systems Biology Approach for Constructing Microglial Regulatory Networks of AD

The integrative network analysis pipeline for elucidating the microglial network model, KDs and drug repurposing (Figure 1) is centered on the data-driven reverse-engineering of causal predictive network models of AD [10-12]. This pipeline can then be directly interrogated to not only delineate biological components causally associated with AD, but also to identify KD genes of individual components underlying AD, leading us towards precision drug repurposing.

**Figure 1.**
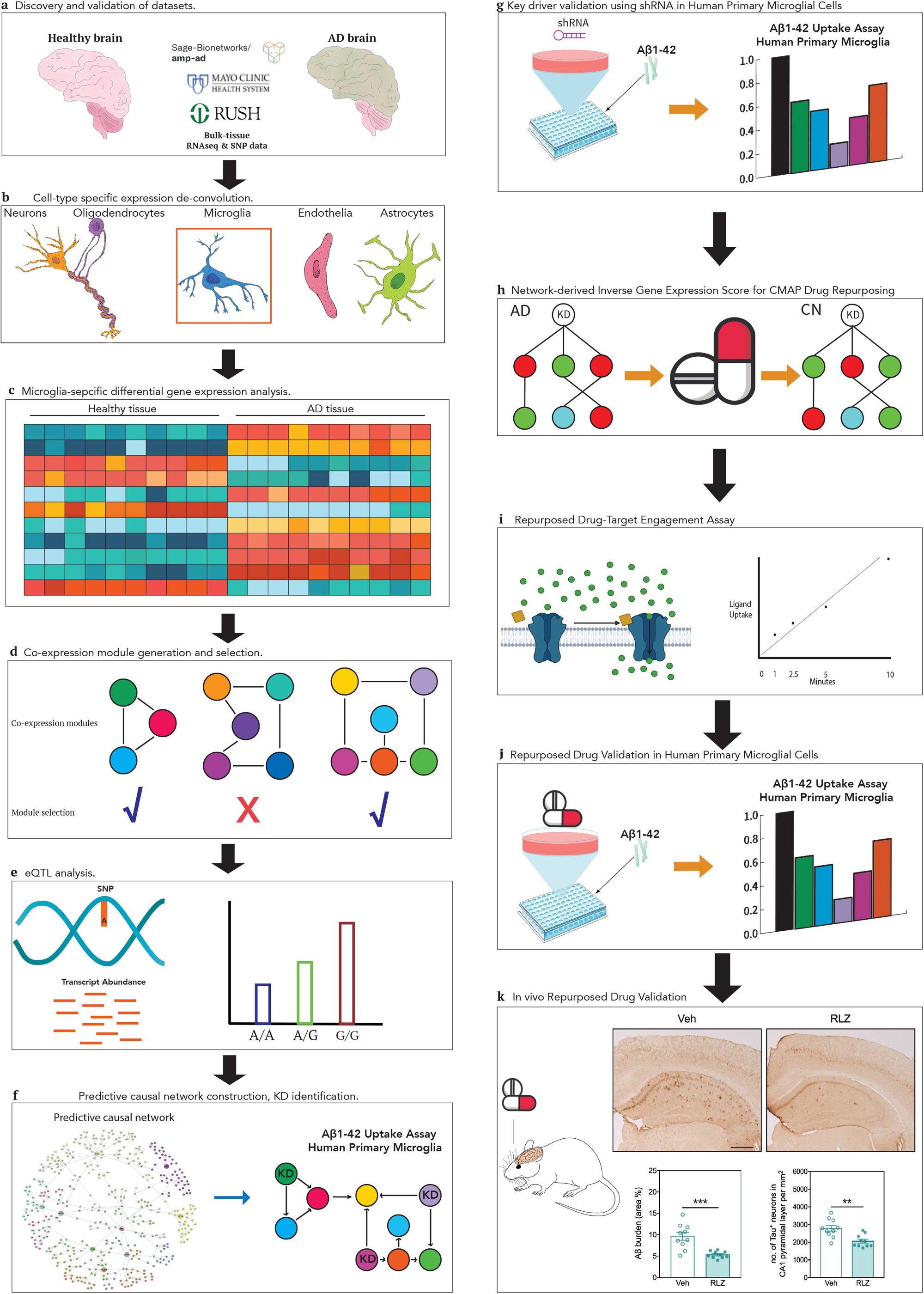
The integrative network analysis pipeline for elucidating the microglial network model, key drivers and drug repurposing for AD with in-vitro and in-vivo validation experiments.

Our pipeline starts with de-convolution of bulk-tissue (post-mortem AD and control brain) RNAseq data into microglial-specific expression residuals with the Population-specific Expression Analysis (PSEA)[13] in ROSMAP and MAYO cohorts, respectively (Step 1, Figure S1). We demonstrated this method is robust against random selection of microglia biomarkers derived from single-cell RNA-seq (scRNA-seq) studies [14-18](Online Method 5). Given the microglial expression residuals in each cohort, we next focused the input of molecular traits into the network reconstruction algorithm on those traits associated with AD, by deriving microglial-specific gene differential expression (DE) signatures (Step 2, Figure S1, Online Method 6.2). To identify co-regulated gene expression traits associated with AD, we constructed gene co-expression networks (Online Method 6.3), and from these networks pinpointed highly correlated sets of genes (modules) that were greatly enriched for DE signatures associated with AD and cell type biomarkers derived from scRNA-seq studies. Each module is then characterized by significantly enriched pathways (Step 3, Figure S1, Online Method 6.8). To derive a final set of genes for seeding the causal predictive network construction process in each cohort, we merged genes in the selected subset of co-expression modules in each cohort respectively (the seed set, Step 5, Figure S1, Online Method 6.4). Given our microglia-centered input set of gene expression traits for the network constructions in AD, we mapped expression quantitative trait loci (eQTLs) for microglial-specific gene expression traits to incorporate the QTLs as structure priors in the network reconstructions, given they provide a systematic perturbation source that can boost the power to infer causal relationships (Step 4, Figure S1, Online Method 6.1)[8, 19]. The input gene set, and eQTL data from MAYO and ROSMAP were then processed by the predictive network to construct probabilistic causal networks of AD in each cohort respectively (Step 6, Figure S1, Online Method 6.7). We next applied a statistical algorithm to detect KD genes in each given network structure[20] and to identify and prioritize master regulators in the AD networks of each cohort respectively (Figure S1, Step 7, Online Method 6.5). These KDs were then prioritized based on ranking score (Online Method 6.6) that resulted in a final set of 9 top-prioritized KDs shared by both cohorts for which we performed functional validation in primary cultures of human microglia. Next, we developed a network-driven inverse gene expression score (Online Method 7, Figure S1, Step 8) to repurpose drugs in the CMAP database against *SLC1A2* (one of the top 9 KDs) in each cohort respectively. To derive robust results, it is essential to point out that we performed every step of our analysis in the two cohorts independently. We then agnostically identified microglial-specific KDs associated with AD and repurposed drugs in each cohort separately. At every step, we cross-validated the results from the two independent analyses. The entire analysis workflow for the independent datasets, resulting in this final group of replicated targets, is illustrated in Figure 1.

### Identifying AD-associated gene signature in microglia and mapping their eQTL

To identify AD-associated signature of microglia gene expression, we performed DE analysis in the ROSMAP and MAYO cohorts using the deconvoluted microglial-specific expression residuals (Online Method 5.1). By comparing expression data between AD and pathologically confirmed controls (CN), there were 513 significant DE microglia-specific genes in the MAYO dataset (hereby “MAYO-microglial”) and 1,693 significant microglial-specific genes in the ROSMAP dataset (hereby “ROSMAP-microglial”) (Figure 2a,b, FDR<0.05), with 120 significant DE genes overlapping between the two sets (Table S1, Fisher Exact Test, odd ratio=3.8169, p-value<2.2E-16). Thus, the independently derived MAYO-microglial and ROSMAP-microglial DE signatures cross-validated each other. We performed pathway enrichment analysis (Online Method 6.8) on the two microglial-specific DE signatures respectively using Human ConsensusPathDB (CPDB)[21] and identified 156 and 365 significantly enriched pathways (Figure 2c) in MAYO and ROSMAP cohort respectively, with 24 pathways significantly dysregulated in both cohorts (Table S2, p-value<0.05), including focal adhesion[22], amino acid and lipid metabolism[23], NF-kB activation[24], Fc gamma R-mediated phagocytosis and phagosome[25], some of which have previously been implicated with AD. By comparing our microglial-specific DE signatures derived from deconvoluted microglial gene expression to the scRNAseq-derived microglial signatures in AD (Online Method 5.3), we demonstrated that our deconvoluted microglial gene expression captured activated microglial states in response to AD pathology.

**Figure 2.**
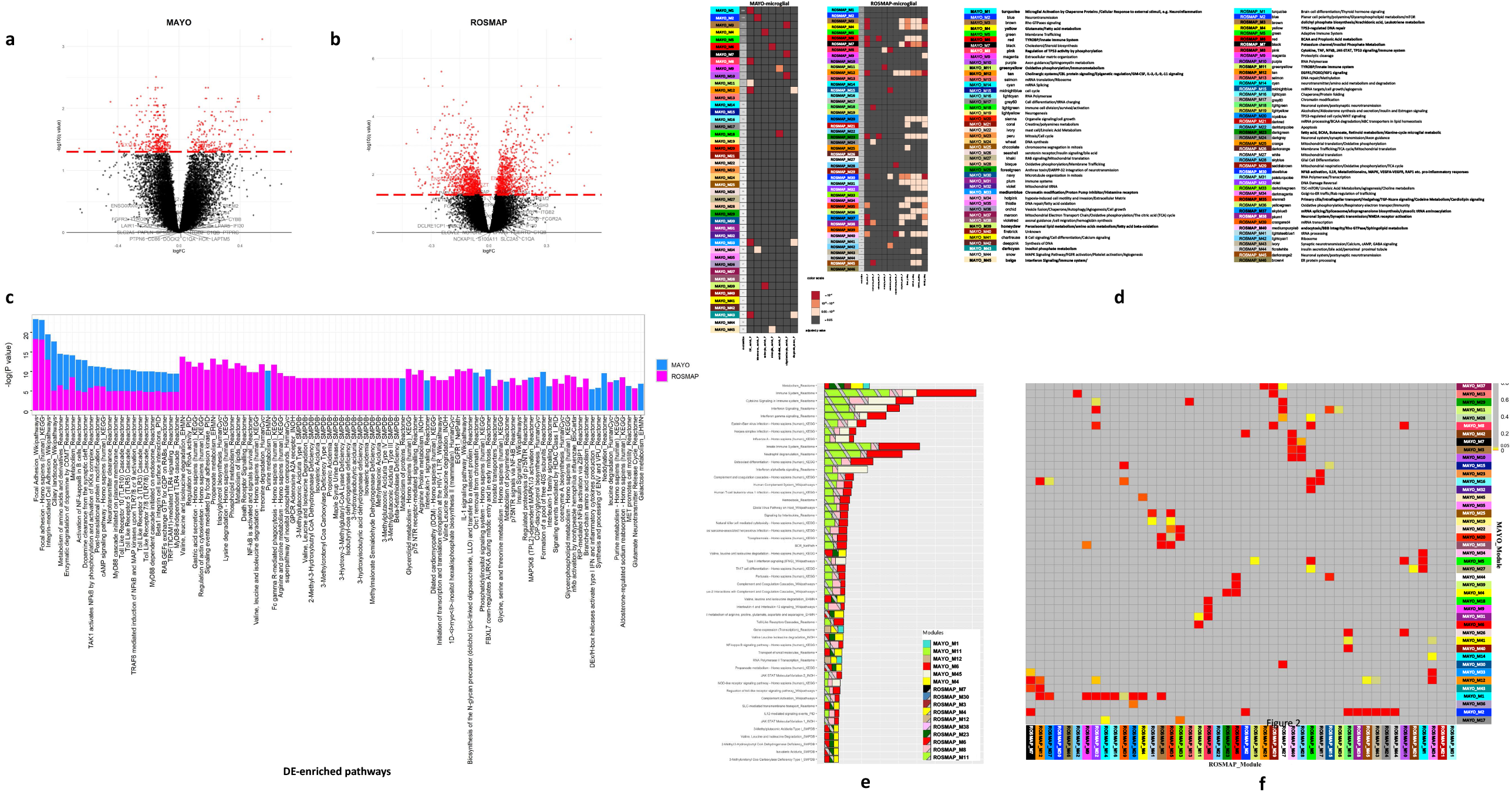
Microglial-specific differential expressed gene signature, co-expression modules and pathway associated with AD. (a) microglial-specific DE gene signature of AD in MAYO; (b) microglial-specific DE gene signature of AD in ROSMAP; (c) Significantly enriched pathways by microglial-specific DE gene signature in each cohort respectively; (d) characterization and functional relevance of microglial-specific gene modules to AD in MAYO and ROSMAP cohort respectively. (e) Pathways significantly enriched by selected microglial-specific gene modules associated with AD in each cohort respectively. (f) Module overlapping heatmap identified 111 pairs of micorglial module with significant overlap.

In addition to the DE signature, another critical input for construction of predictive network models, are expression quantitative trait loci (eQTL). We mapped *cis*-eQTL by examining association of microglial-specific expression residuals with genome-wide genotypes assayed in the ROSMAP and MAYO cohorts, respectively (Online Method 6.1). In the MAYO-microglial and ROSMAP-microglial, 3875 (19.5%) and 5186 (25.6%) genes were significantly correlated with allele dosage (FDR < 0.01, Table S4). Of the *cis*-eQTL detected in each cohort, 1785 genes were overlapping between the two sets (46% of MAYO *cis*-eQTLs and 34% of ROSMAP cis-eQTLs, Fisher’s Exact Test p-value < 2.2e-16), further suggesting that independent analysis of the two cohorts cross-validated each other.

### Identifying gene co-expression modules of microglia associated with AD

To further identify co-regulated set of genes (modules) that are likely to be involved in common biological processes under AD as input gene set for the predictive network, we clustered the microglial gene expression traits into data-driven co-expression networks (Online Method 6.3). We constructed co-expression networks on the deconvoluted microglial residuals of AD samples within each dataset, resulting in MAYO-microglial co-expression network consists of 45 modules ranging in size from 32 to 2,201 gene members and the ROSMAP-microglial co-expression network consists of 46 modules ranging in size from 115 to 892 gene members (Figure 2d). To evaluate functional relevance of each microglial-specific modules to AD pathology, we performed enrichment analysis of each module for its AD-associated microglial DE signatures and known single-cell marker genes for the 5 major cell types in the CNS[16], and categories of AD traits available from its respective cohort (Figure 2d). Based on the enrichment results, the final set of 10 and 13 microglial-specific modules were selected for causal network modeling of AD in each cohort, respectively (Online Method 6.4).

To characterize mechanisms involved in the co-expression modules, we performed pathway enrichment analysis to identify biological processes across all modules in each cohort (Table S5). Out of the selected modules, we identified that 373 and 481 CPDB pathways were significantly enriched by 6 out of 10 and 10 out of 13 selected modules from the MAYO and ROSMAP co-expression networks respectively, with 241 pathways overlapping between the two datasets (Figure 2e, FDR<0.05, Fisher’s Exact Test, OR=30.11, p-value<2.2E-16, Table S5). These pathways, such as microglia pathogen phagocytosis, interferon signaling and immune response, microglial activation mediated by chaperone proteins, TP53-regulated DNA repair and NF-kB activation, highlighted key microglial-specific processes associated with AD. In comparing all modules between the datasets, we identified 111 module pairs with significant overlap (Figure 2f, FDR<0.05) demonstrating the robustness of two independent co-expression networks.

### Predictive Network Modeling of Genetic Regulations Identified Pathological Pathways and Key Drivers for Microglial Function in AD

To build causal network and identify microglial-specific KDs, we employed our cutting-edge predictive network modeling pipeline [10-12] on the set of seeding genes which are pooled from the 10 and 13 selected modules respectively in each cohort (4187/5152 genes for MAYO/ROSMAP, Online Method 6.7). The overlap between the two seeding gene sets contains 1842 genes (35.7% of ROSMAP and 43.9% of MAYO; Fisher’s Exact Test, p-value <2.2e-16), indicating reproducibility of these analyses across these two independent datasets. Therefore, analysis using these two datasets increases the power to discover high-confidence microglial KDs that are associated with AD pathology. We also incorporated *cis*-eQTL genes into each network construction as structural priors in the ROSMAP and MAYO dataset respectively. Of particular note, the 5186 and 3875 unique *cis*-eQTL genes identified in the ROSMAP-and MAYO-specific datasets, 1978 and 687 cis-eQTL genes overlapped with the seeding genes in each network respectively (Figure 3a,b).

**Figure 3.**
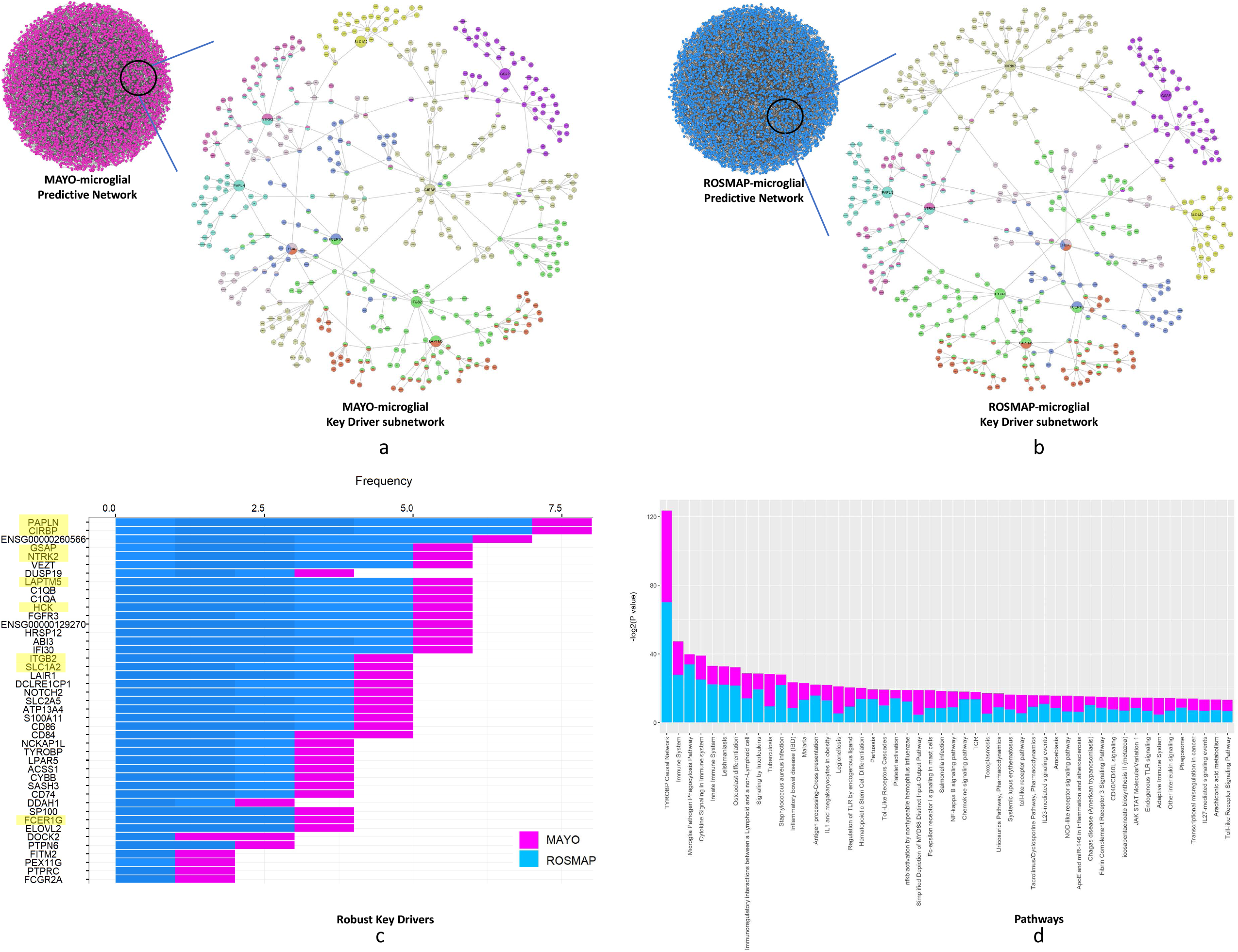
Predictive network models of causal genetic regulations identified key drivers for microglial function in AD. (a) predictive network model of microglial functions associated with AD in MAYO cohort (Left: hairball containing XXX causal genetic regulations among 4187 genes, Right: a zoom-in subnetwork within 3-step downstream of the key drivers for microglial functions in AD). (b) predictive network model of microglial functions associated with AD in ROSMAP cohort (Left: hairball containing XXX causal genetic regulations among 5152 genes, Right: a zoom-in subnetwork within 3-step downstream of the key drivers for microglial functions in AD). (c) 43 key drivers replicated across both bohorts modulating microglial network states in AD. (d) Functional characterization of the 43 KDs by pathway enrichment analysis on the downstream sub-network of these KDs in the ROSMAP-and MAYO-microglial networks respectively, revealed they are robust upstream regulators of microglial phagocytosis process associated with AD.

To identify KD genes that modulate network states under AD, we applied Key Driver Analysis [20] and identified 757 and 164 KD genes in the ROSMAP-and MAYO-microglial networks respectively with 43 KDs replicated across both networks (Figure 3c). To characterize biological functions of the replicated 43 KDs, we performed CPDB pathway enrichment analysis on the downstream sub-network of these KDs in the ROSMAP-and MAYO-microglial networks respectively, which revealed that both sub-networks are significantly (p-value<0.05) enriched for phagocytosis and phagosome pathways and these 43 KDs are robust upstream regulators of phagocytosis which is a microglial-specific process associated with AD (Figure 3d, Table S6).

### Validation of key drivers as master regulators of microglia function associated with AD

To validate microglia-mediated phagocytic function of these 43 KDs associated with AD, we prioritized these genes (Online Method 6.6) and selected a total of 9 novel KDs, consisting of top four prioritized KDs with frequency between 5 to 7.5 (*PAPLN, CIRBP, GSAP, NTRK2*) and the top two with frequency below 5 (*ITGB2* and *SLC1A2*). It is noteworthy that functions of these genes have not yet been associated with microglial phagocytosis. We utilized three additional KDs (i.e., *LAPTM5, HCK, FCER1G*) that have been previously implicated in phagocytosis [26-28] as positive controls. To further confirm our assay, we also included knockdown of TREM2 as well as treatment with cytochalasin D and ice-cold temperature (i.e., 4°C) as control conditions that are known to regulate phagocytotic function in microglia [29, 30].

We analyzed uptake of fluorescently labelled Aβ1-42 via confocal microscopy in human primary microglia (HPM) that were transfected with lentiviral clones containing unique short hairpin RNA (shRNA) targeting one of the nine selected KDs, which were then compared to HPMs from the same donor that were transfected with a control (i.e., non-targeting) vehicle (Online Method 7, 8, 9.1). Our results showed that genetic knockdown of all nine prioritized KDs resulted in a significant reduction in cytotoxic Aβ1-42 uptake by HPM as compared to the vehicle control. These data imply a critical role for these KDs in regulating microglial phagocytosis under AD conditions (Figure 4, Table S7). As expected, all positive controls resulted in a significant decrease in Aβ1-42 uptake by HPMs.

**Figure 4.**
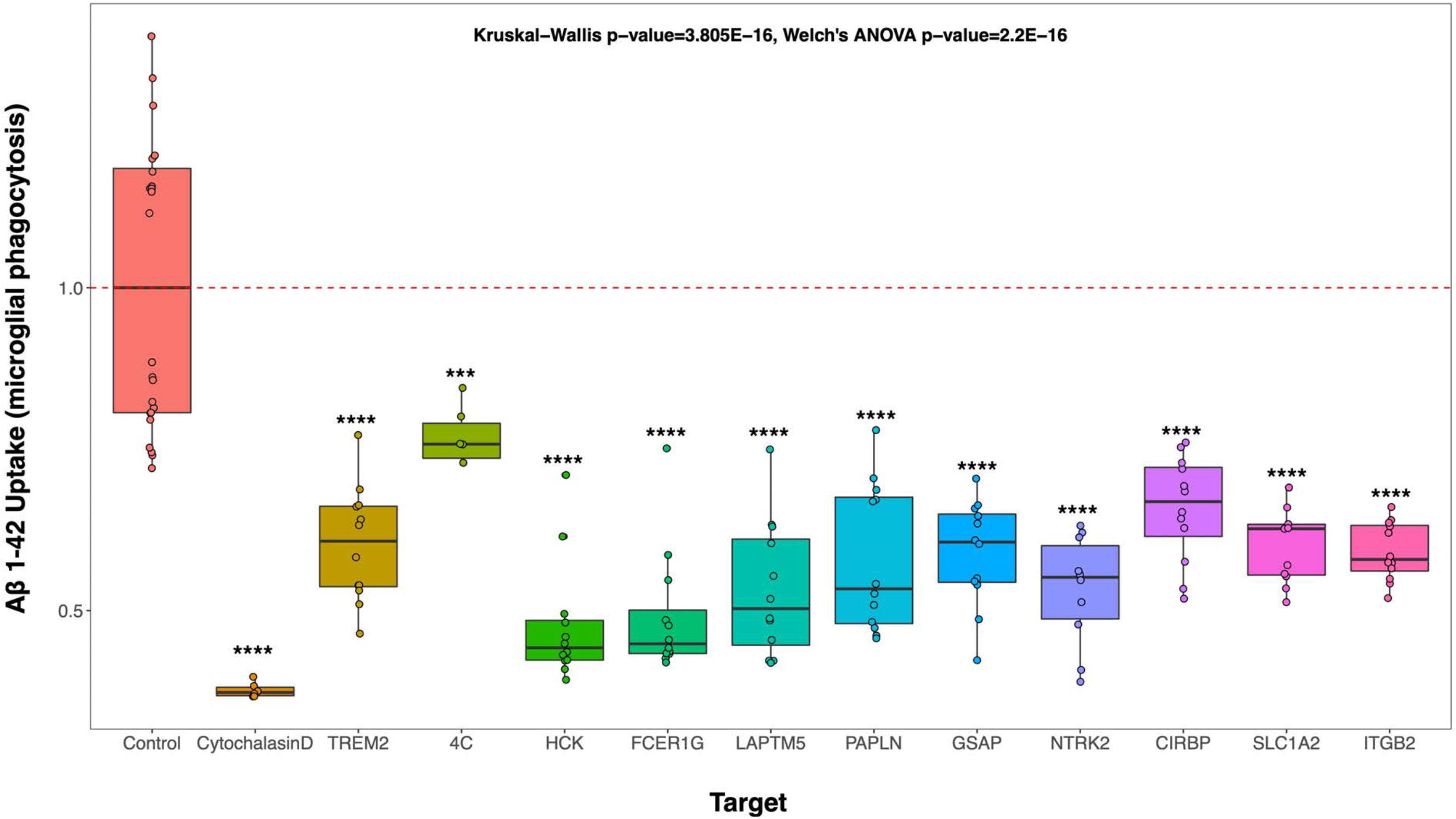
Knockdown of all 9 prioritized key drivers for microglial-mediated phagocytosis resulted in a significant reduction in cytotoxic Aβ1-42 uptake by human primary microglial cells as compared to the vehicle control under AD condition. Positive controls: Cytochalasin D, 4C condition, knowck down of *LAPTM5, HCK, FCER1G* and *TREM2* which have been implicated in regulation of microglial phagocytosis.

### Drug repurposing against key drivers as potential treatments for AD

To further demonstrate the therapeutic potential of our validated KDs, we repurposed drugs from the CMAP/LINCS database [31] against each of the validated nine KDs (Online method 7). In the following illustration, we highlighted repurposing against *SLC1A2*, where we identified a total of 26,375 and 59,519 significant perturbation hits (out of total 720,216 perturbations in CMAP) in MAYO and ROSMAP cohorts, respectively, with 11,650 overlapped perturbations over 207 cell lines. Out of these perturbations, 1425 and 762 significant (FDR < 0.05) perturbation hits are derived from six CNS cell lines in CMAP for MAYO and ROSMAP cohorts, respectively, with 292 hits overlapped. These 292 overlapped perturbations are mapped to 228 (out of total 33,609) drugs in CMAP, indicating their robust reversal effects on downstream gene expression of *SLC1A2* in ROSMAP-and MAYO-microglial network (Figure 5a). We focused on validating the effects of riluzole, one of the top 228 significant repurposed drugs against *SLC1A2*, on microglial functions associated with AD.

**Figure 5.**
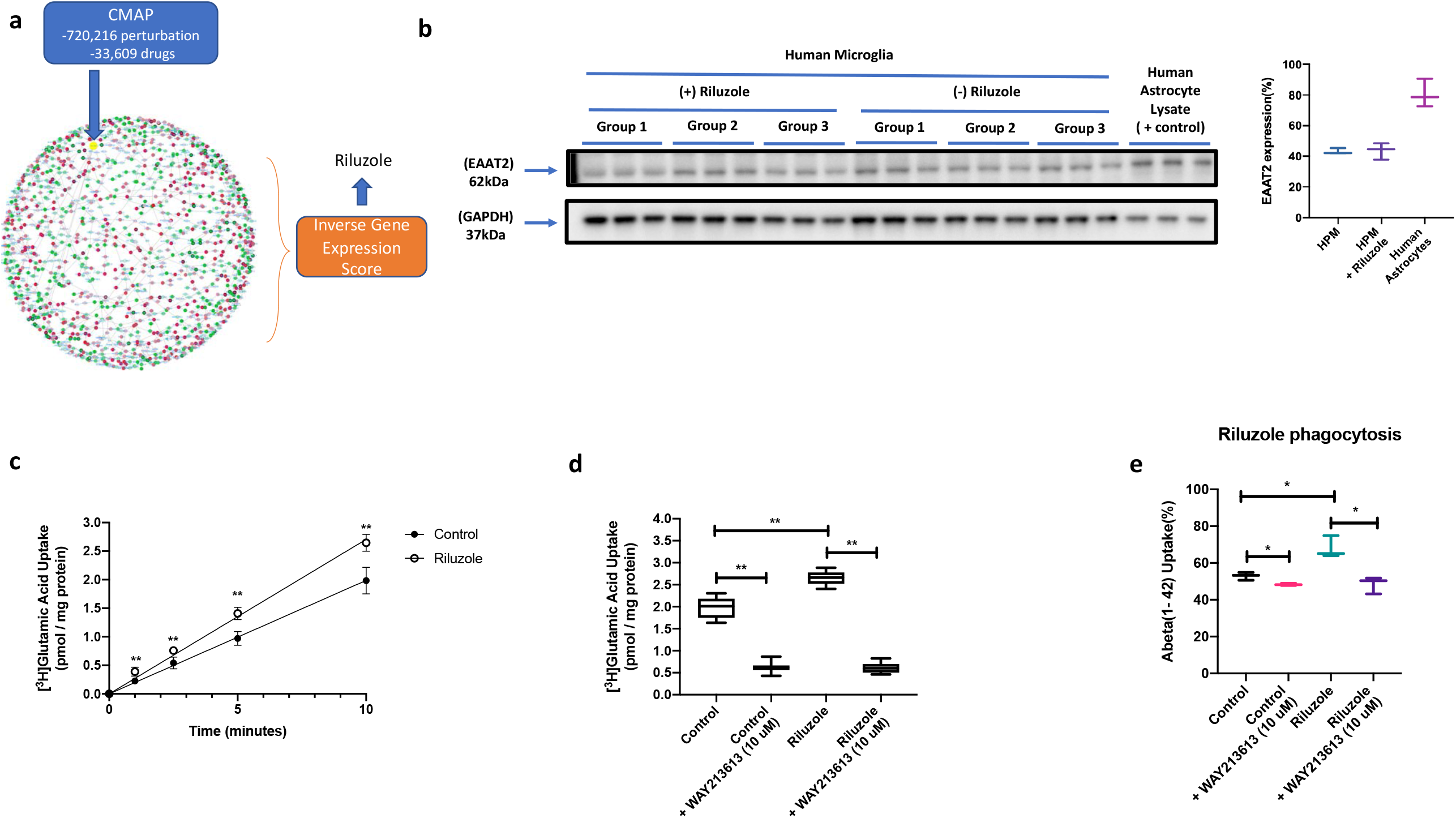
Drug repurposing in CMAP database against *SLC1A2* (one of the 9 key drivers for microglial phagoctysis in AD) and in vitro validation of riluzole effects on *SLC1A2* functional expression and Aβ1-42 phagocytosis in HPMs. (a) Illustration of drug repurposing in CMAP database with Network-driven Inverse Gene Expression Score against *SLC1A2*, where we identified 228 significant drugs in CMAP showing reversal effects on downstream gene expression of *SLC1A2* in ROSMAP-and MAYO-microglial network. We focused on validating the effects of riluzole, one of the top 228 significant repurposed drugs against *SLC1A2*, on microglial functions associated with AD. (b) *SLC1A2* protein expression was measured by western blot analysis of HPMs in the presence and absence of riluzole. Cell membrane preparations were resolved on a 4-12% SDS-polyacylamide gel, transferred to a polyvinylidene difluoride membrane and analyzed for expression of *SLC1A2* or *GAPDH* (i.e., the loading control). Each lane triplet on the depicted western blot corresponds to a HPM sample derived from a single treatment. The image shows a representative blot from three separate experiments (n = 3). Relative levels of *SLC1A2* protein expression were determined by densitometric analysis and normalized to *GAPDH* (c) The rate of uptake of [^3^H]glutamic acid, a prototypical *SLC1A2* transport substrate, was measured in HPMs in the presence and absence of riluzole. (d) Cellular accumulation of [^3^H]glutamic acidwas measured in control and riluzole-treated HPMs in the presence and absence of WAY213613 (10 μM), an established competitive *SLC1A2* inhibitor. (e) Uptake of fluorescent Aβ1-42 in control and riluzole-treated HPMs *in* the presence and absence of WAY213613 (10 μM). Cellular accumulation of Aβ1-42 is an indicator of phagocytosis. Quantitative data are expressed as mean ± SD from three independent experiments with each data point in an individual experiment representing 3-4 measurements. Asterisks indicate data points that are statistically significant from control (* p < 0.05; ** p < 0.01).

### Novel function of riluzole in activation of microglial phagocytosis in AD

Riluzole is an FDA-approved neuroprotective drug for treatment of amyotrophic lateral sclerosis (ALS) [32]. Studies in primary cultures of mouse astrocytes showed that riluzole increases protein expression and transporter activity of *SLC1A2* (also known as excitatory amino acid transporter 2 (*EAAT2*) or glutamate transporter 1 (*GLT-1*)), thereby facilitating synaptic removal of glutamate and protection of neurons against excitotoxicity [33]. Of particular relevance to our work, *SLC1A2* protein expression has been detected by western blot analysis in mouse [34], rat [35], and human [36] microglia; however, riluzole’s effect on *SLC1A2*-mediated microglial functions, and specifically microglial phagocytosis associated with AD, has not been previously evaluated. Therefore, we performed i) transport assays to measure effects of riluzole on *SLC1A2*-mediated uptake of glutamic acid; and ii) fluorescent Aβ1-42 uptake experiments to link *SLC1A2*-mediated transport activity to regulation of microglial phagocytosis. Protein expression of *SLC1A2* was measured by western blot analysis and confirmed that riluzole treatment (10µM for 24h) did not increase *SLC1A2* protein expression in HPMs (Figure 5b); however, [^3^H]glutamic acid (1.0 µCi/ml) uptake was significantly increased in riluzole treated human microglia than in untreated control cells (Figure 5c, p-value<0.01). Transport specificity was further confirmed using the competitive *SLC1A2* inhibitor WAY213613 (10µM), which significantly reduced [^3^H]glutamic acid uptake (p-value<0.01) in both control and riluzole-treated HPMs (Figure 5d). Furthermore, our experiment showed phagocytosis of Aβ1-42 was significantly increased in HPMs treated with riluzole (Figure 5e, p-value < 0.05) as compared to untreated control cells. The *SLC1A2* inhibitor WAY213613 significantly (p-value < 0.05) reduced fluorescent Aβ1-42 uptake in both control and riluzole-treated HPMs (Figure 5e). Overall, our in vitro experiments confirmed that riluzole significantly enhanced microglial phagocytosis of Aβ1-42 associated with AD via increasing *SLC1A2*-mediated transport activity but not its protein expression.

### Therapeutic potential of riluzole associated with microglial phagocytosis of amyloid burden and tau pathology in AD

Next, we examined, *in vivo*, the therapeutic potential of riluzole associated with microglial phagocytosis on AD in 16-month-old female 3xTg-AD mice, which exhibit both amyloidosis and tauopathy. Only female mice were used in this component of our study because female 3xTg-AD mice exhibited more stable and severe AD-related pathologies than those present in males [37]. We treated 3xTg-AD mice with daily riluzole injection (4 mg/kg, i.p.) for 4 weeks. Our results showed that riluzole significantly increased expression of *SLC1A2* in the hippocampus of 16-month-old 3xTg-AD mice (p-value < 0.05, Figure 6a). Although riluzole did not affect the hippocampal levels of soluble Aβ1-40 (Figure 6b) or Aβ1-42 (Figure 6c), it significantly reduced amyloid burden (p-value < 0.01, Figure 6d). Additionally, riluzole treatment alleviated tauopathy by significantly decreasing the densities of total Tau and pTau(Thr181)-immunoreactive neurons in the CA1 pyramidal layer of the dorsal hippocampus (Figure 6e). Differences in pTau(AT8)-and pTau(S396)-immunoreactive neurons between two groups were not significant. In summary, our *in vivo* findings indicate that riluzole-induced upregulation of *SLC1A2* and enhancement of microglial phagocytosis decrease amyloid burden and number of p-tau-loaded neurons.

**Figure 6.**
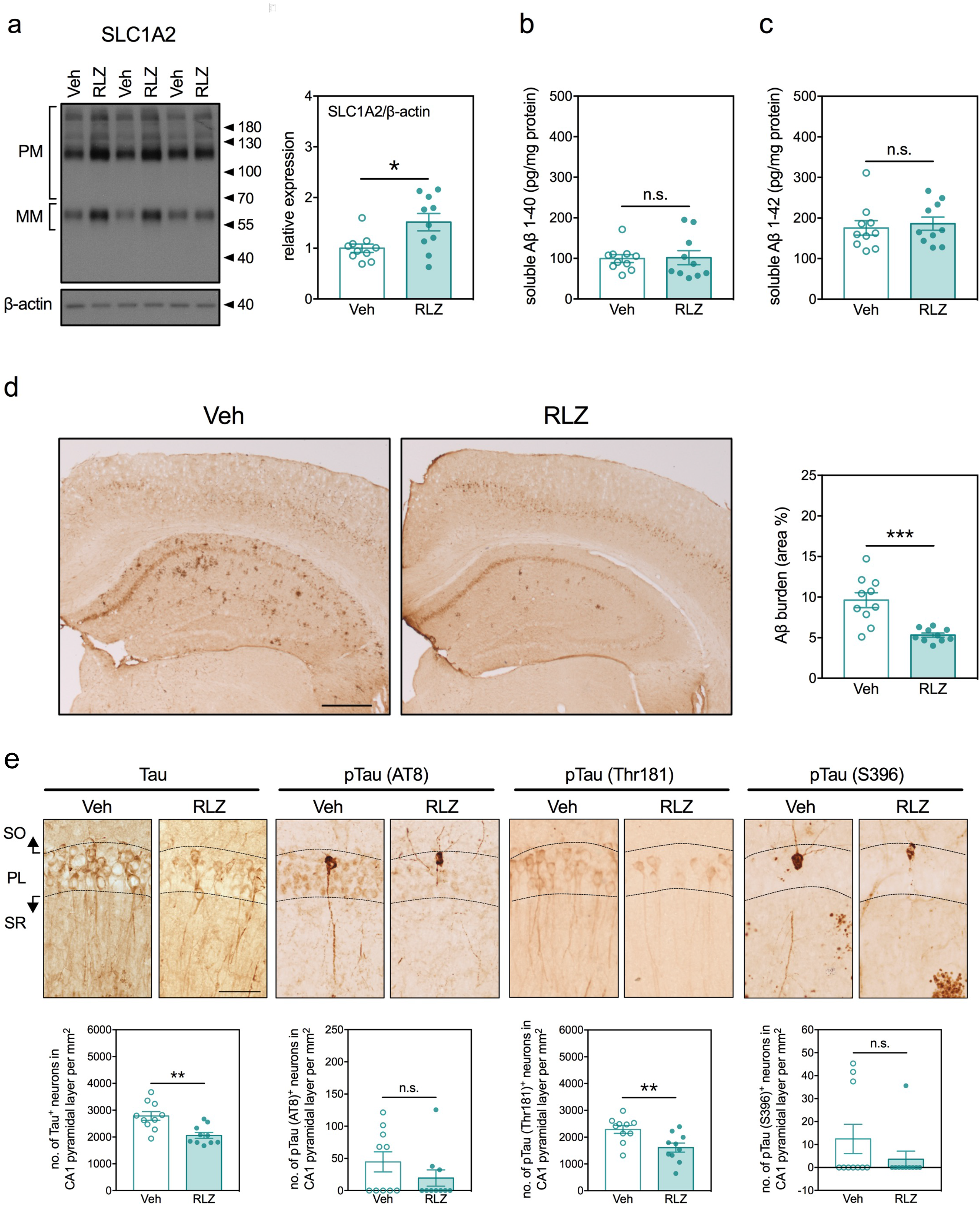
Riluzole upregulated *SLC1A2* expression and reduced amyloidosis and tauopathy in the hippocampi of female 3xTg-AD mice. (a) Representative Western blots (left panel) and quantitative results (right panel) of *SLC1A2* expression in the dorsal hippocampus. (b) Quantitative results of soluble Aβ1-40 in the dorsal hippocampus. (c) Quantitative results of soluble Aβ1-42 in the dorsal hippocampus. (d) Representative micrographs (left panels) and quantitative results (right panel) of amyloid burden in the dorsal hippocampus. (e) Representative micrograph (upper panels) and quantitative results (lower panels) of the densities of Tau+, pTau (AT8)+, pTau (Thr181)+, and pTau (S396)+ neurons in the CA1 region of dorsal hippocampus. SO: stratum oriens, PL: pyramidal neuron layer, SR: stratum radiatum. All data are expressed as mean ± SD. n = 10 mice per group. n.s.: not significant. * p < 0.05, ** p < 0.01, *** p < 0.001.

## Discussion

GWAS studies in LOAD have identified several microglial specific genes; however, it is unclear how these genes are involved in LOAD neuropathology. Hence, a comprehensive characterization of gene regulatory networks of microglia associated with the disease is critical to provide insights into underlying mechanisms of AD. Our study used an innovative systems biology approach to build causal network models of microglial component of postmortem human brain samples with AD and to identify microglia-specific KDs and cellular pathways that contribute to AD pathology using the MAYO and ROSMAP datasets in the AMP-AD consortium. We first deconvolved RNA-seq data from region-specific brain tissue in MAYO (TCX-Temporal Cortex, 79 AD and 76 CN) and ROSMAP (PFC-Pre-frontal Cortex, 212 AD and 200 CN) cohorts into microglial-specific gene expression component. Next, we derived microglial-specific differential expression signature, cis-eQTLs, co-expression networks, predictive causal network models and KDs in each dataset respectively. We successfully validated 9 novel KD genes associated with microglial phagocytosis in AD, specifically *HCK, FCER1G, LAPTM5, ITGB2, SLC1A2, PAPLN, GSAP, NTRK2 and CIRBP*. The robust pathways enriched by these networks and key driver genes confirmed phagocytosis is one of the central biological processes for microglia under AD pathophysiological conditions.

Indeed, microglial phagocytosis plays a critical role in AD pathophysiology, particularly with respect to amyloid and hyper-phosphorylated tau protein clearance[5]. To date, much of the research on molecular mechanisms associated with microglial phagocyotsis has focused on the role of triggering receptor expressed on myeloid cells (*TREM1 and 2*) because Aβ oligomers and *APOEs* are ligands for these receptors [38, 39]. Many of the upstream and downstream regulators that drive microglial phagocytosis independently or in conjunction with *TREM1* and *TREM2* have not been rigorously validated. Our network models illustrated that *LAPTM5, ITGB2* and *TREM2* co-regulate each other through a feedback loop, while *GASP, FCER1G, SLC1A2, HCK, PAPLN, CRIBP* are down-stream effectors of *TREM2* (Online Method 6.9, Supplementary Figure S6), although further functional validation experiments are required to confirm these associations. In fact, the phagosome pathway is activated upon tissue injury demarcated by neuronal loss or amyloid plaque buildup, two important pathophysiological hallmarks of AD. It is known that microglia colocalize with amyloid beta plaques, suggesting neuroprotective functions in regulating plaque burden and slowing AD progression. In contrast, microglia can trigger neuronal death by phagocytizing “stressed-but-viable” neurons, which can contribute to further neuronal loss [40, 41]. Therefore, fine-tuning microglia phagocytosis in AD represents a promising therapeutic strategy.

Among our KDs of microglial phagocytosis, Castillo and colleagues [42] showed that both HCK and FCER1G are upregulated in the cortex of *App*^NL-G-F/NL-G-F^ transgenic mice during progression of amyloidosis. *HCK* is a member of the Src family of protein tyrosine kinases that couples to Fc receptors during phagocytosis [43, 44]. *FCER1G* encodes for a IgE Fc receptor subunit and has been implicated as a hub gene for microglial activity both in amyloid overexpressing models and in a LOAD transcriptomics study [8, 45]. *LAPTM5* is associated with lysosomal organization and biogenesis[46, 47]. A recent study using a murine amyloid responsive network and GWAS defined association has shown that genetic variations in *LAPTM5* are associated with amyloid deposition in AD [48]. While it is tempting to associate *LAPTM5* with phagocytosis and synaptic remodeling in the AD brain, its direct function in microglia phagocytosis requires further validation [26]. Our study demonstrated for the first time that this gene directly regulates microglial phagocytosis of Aβ1-42. *NTRK2* encodes the tyrosine kinase receptor B (*TRKB*), the receptor for brain-derived nerve growth factor (BDNF). Indeed, established neurotrophic properties of BDNF point towards a central role for *TRKB* in promoting neural repair in response to pathophysiological stressors in the AD brain [49]. A reduction of BDNF-TRKB signaling in microglia elicit their activation and neurotoxic effect [50]. Additionally, kinase activity of *TRKB* functions in dephosphorylation of hyper-phosphorylated tau protein [51].

*GSAP* forms complexes with γ-secretase, which cleaves amyloid β-protein precursor (APP) to form Aβ. Whereas clinical trials on direct γ-secretase inhibitors have been unsuccessful, *GSAP* remains a potential therapeutic target [52]. Specifically, *GSAP* controls both γ-secretase substrate specificity and enzymatic activity[53], thereby providing an opportunity to improve target engagement in the APP cleavage pathway. *CIRBP* is a constitutively expressed cold-shock protein that stabilizes target transcripts in the event of mild-cold stress in mammals [54]. In contrast to *GSAP*, the antioxidative and antiapoptotic mechanisms of *CIRBP* are known to reduce Aβ toxicity [55]. As a member of the solute carrier family that is preferentially expressed in astrocytes, *SLC1A2* functions in removal of glutamate from the synapse and reduced function of this transporter can lead to excitotoxicity [56]; however, *SLC1A2* function in microglia has not previously been evaluated. We report, for the first time, that this gene regulates microglial phagocytosis of Aβ1-42. *ITGB2* encodes the immunomodulatory protein CD18, which has a role in leukocyte adhesion [57]. *PAPLN* exhibits peptidase and serine-type endopeptidase inhibitor activity [58]. As compared with our other KDs, no published data on phagocytosis-related functions such as docking or cellular uptake are available in the scientific literature for PAPLN.

We also developed a network-driven inverse gene expression score to repurpose drugs in CMAP database against novel targets for AD treatment. and predicted that riluzole could potentially affect microglial phagocytosis. This hypothesis was validated by our experiments using both *in vitro* cell culture systems and *in vivo* AD models. Using HPMs, we demonstrated that *SLC1A2* transport activity is required to stimulate phagocytosis as evidenced by reduced uptake of fluorescent Aβ1-42 in cells treated with WAY213613. *In* vivo, we showed that riluzole decreased amyloid plaque load without affecting soluble Aβ1-40 and 1-42 levels. Pools of soluble Aβ are dynamically affected by synthesis, clearance, and aggregation/deposition. Several mechanisms are known to mediate clearance of brain Aβ, including receptor-mediated and/or transporter-mediated Aβ efflux from the brain, direct drainage to the cervical lymph nodes via the meningeal lymphatic system, and enzymatic and phagocytic degradation. Our findings indicate that riluzole-induced upregulation of *SLC1A2* and enhancement of microglia-mediated Aβ clearance shifted the equilibrium from insoluble pools of Aβ to soluble pools of Aβ resulted in decreases in Aβ deposition. The reduced amyloid burden may also result from direct phagocytosis of aggregated Aβ fibers/protofibrils. In the brain, *SLC1A2* is predominantly expressed in astrocytes that function, in part, to maintain cerebral glutamate homeostasis. The decreased amyloid plaque load may also be attributed to astrocyte-mediated glutamate clearance and reduced excitotoxicity, which collectively with microglial phagocytosis meliorated the degrees of Tau phosphorylation [59]. Nonetheless, our results imply that altered *SLC1A2* protein expression and/or activity in both HPMs and in 3xTg mice contributes to this neuroprotective mechanism mediated by promoting microglial phagocytosis in AD conditions. This observation is supported by a recent phase 2 double-blind, randomized, placebo-controlled study that showed a beneficial effect of riluzole (50 mg twice a day) on brain glucose metabolism and cognitive measures in patients with AD [60].

In summary, our innovative computational systems biology modeling of microglia-specific networks illustrated mechanisms of microglial interactions with AD pathology and identified nine master regulators of microglial phagocytosis in AD conditions: *HCK, FCER1G, LAPTM5, ITGB2, SLC1A2, PAPLN, GSAP, NTRK2* and *CIRBP*. Our in-vitro experiment showed significant reduction in microglial uptake of Aβ1-42 following knockdown of the nine KDs, suggesting they may be potential microglial targets for novel therapeutic developments to design new paradigms for AD treatment. Indeed, as illustrated by our *in vitro* and *in vivo* experiments with riluzole, the repurposed drugs against one of these prioritized targets, *SLC1A2*, offers novel insights for academic and industrial drug discovery programs aimed at developing the next frontier of therapeutics for neurodegenerative diseases such as LOAD.

## Figure Legend

**Figure S1.**
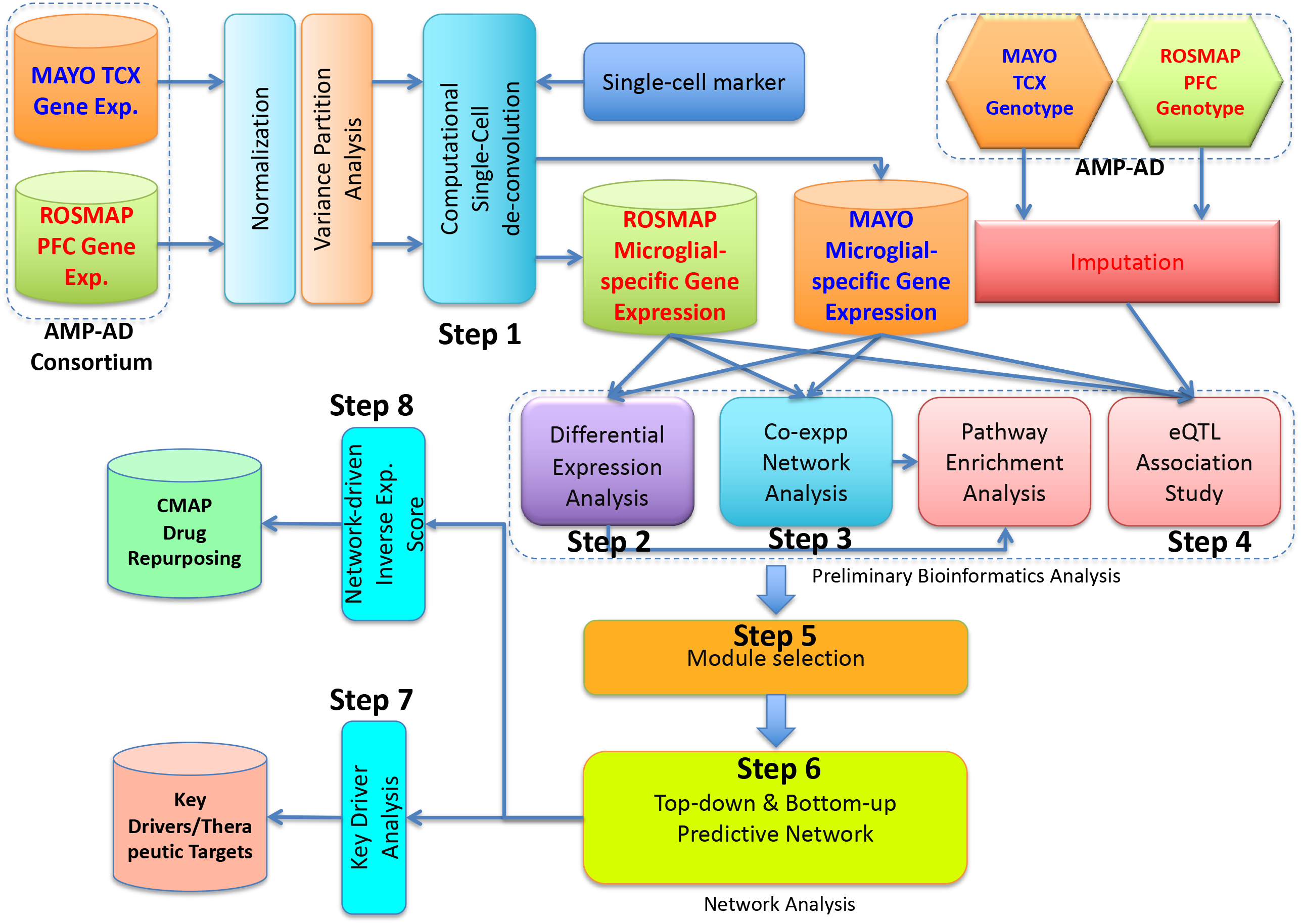
Workflow of our network analysis pipeline integrating two independent brain transcriptome and genome-wide genotype datasets (MAYO and ROSMAP) to construct microglial-specific predictive networks of AD and predict key drivers (therapeutic targets) associated with microglial functions in AD and drug repurposing for AD treatment. TCX: temporal cortex; PFC: prefrontal cortex.

**Figure S2.**
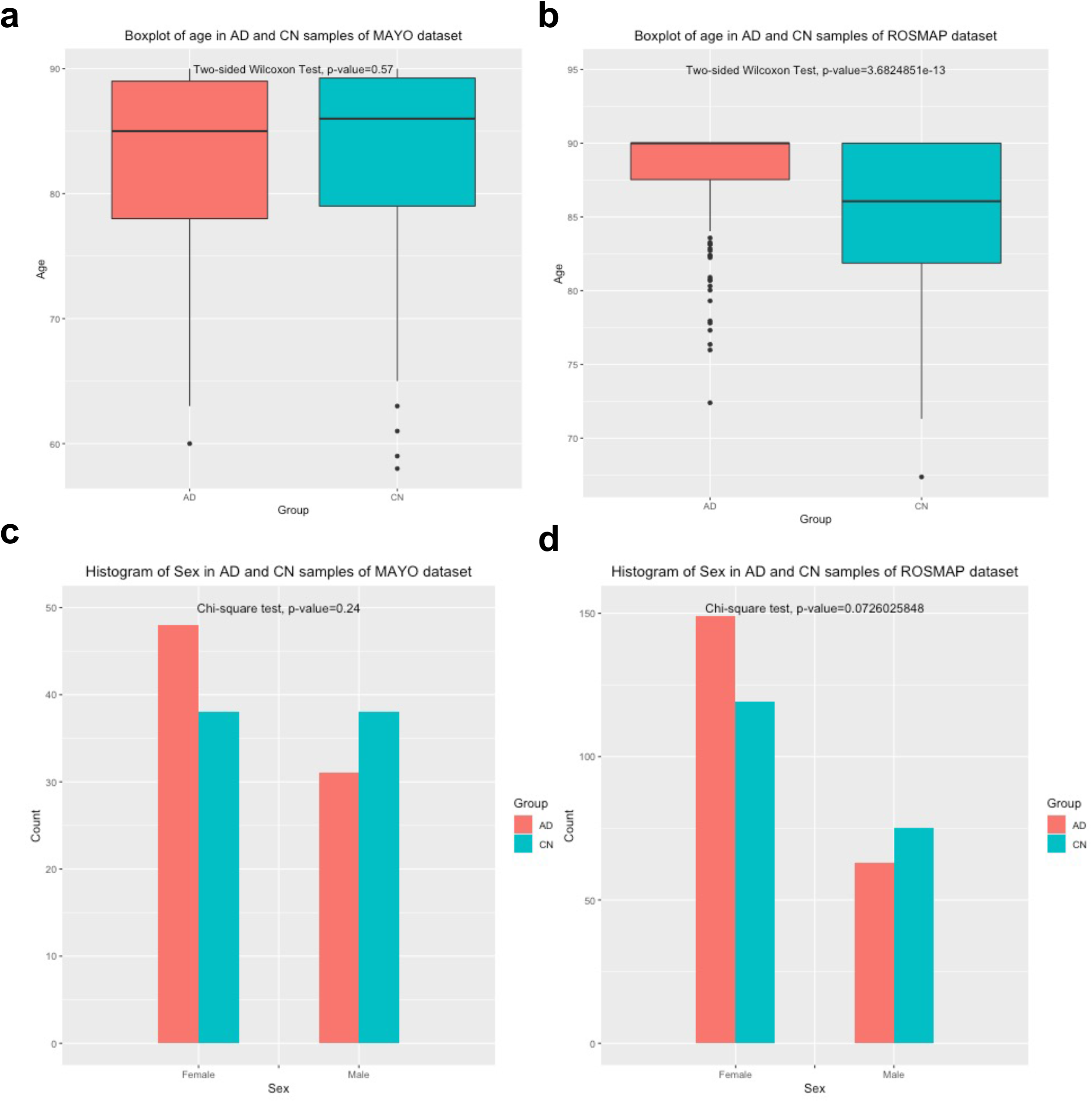
(**a**,**b**) The age distribution of AD and cognitively normal (CN) samples in the MAYO (**a**) and ROSMAP (**b**) datasets, each compared using an unpaired, two-sided Wilcoxon test. There was no significant age difference in MAYO (p-value=0.57) and a significant difference in ROSMAP (p-value=3.68E-13). To remove this effect in ROSMAP, age was adjusted along with other covariates in the ROSMAP residuals. (**c**,**d**) The sex distribution of AD and CN samples in the MAYO (**c**) and ROSMAP (**d**) datasets. A Chi-square test showed no significant difference in the sex breakdown in the MAYO (p-value=0.24) or ROSMAP (p-value=0.0726) datasets.

**Figure S3.**
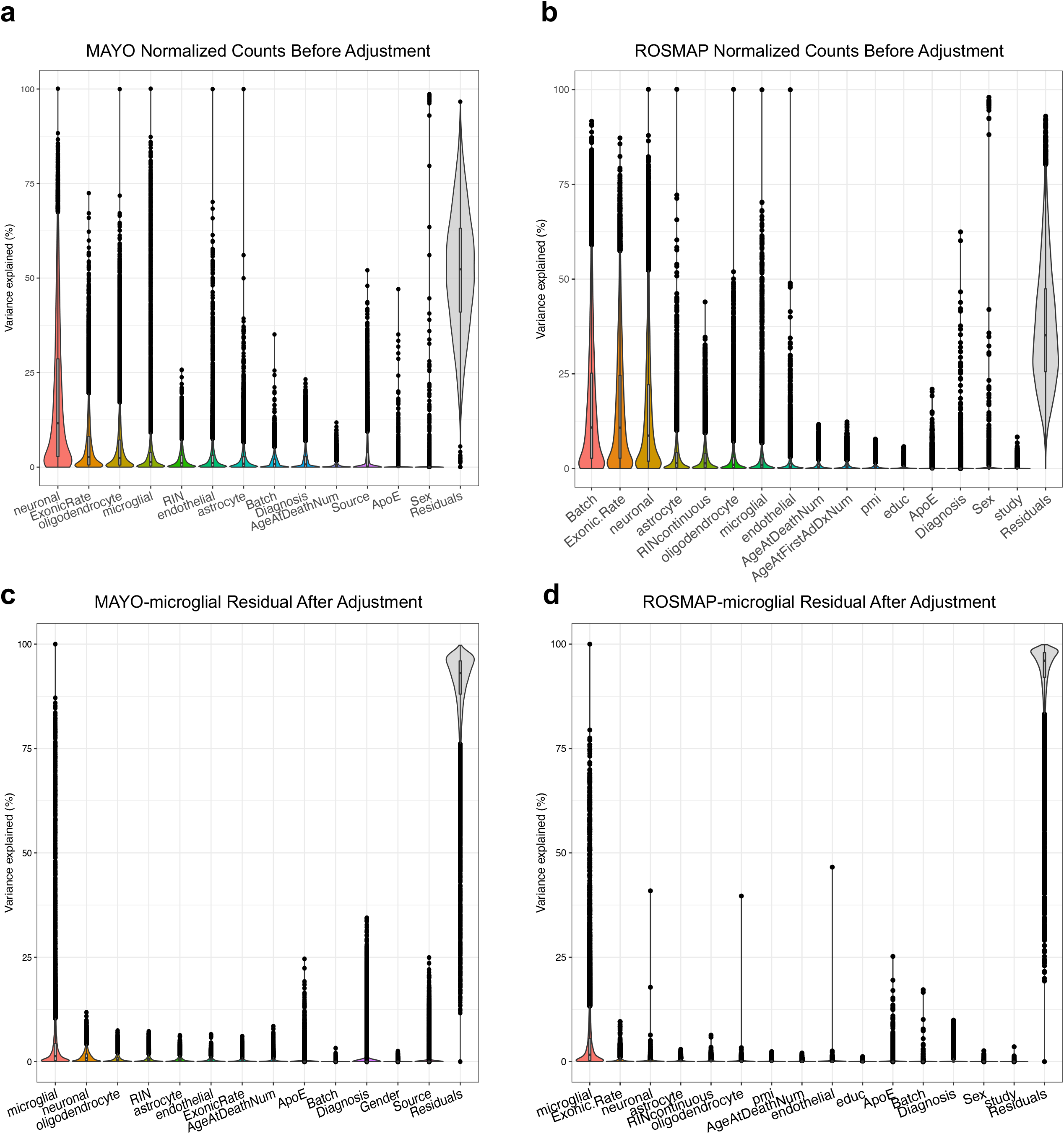
(**a**,**b**) Gene expression variance partition analysis (VPA) of bulk-tissue RNAseq data in the MAYO (**a**) and ROSMAP (**b**) cohorts *<u>before</u>* deconvolution and covariate adjustment reveals a prominent effect of cell type on bulk-tissue gene expression in the brain. *ENO2, CD68, GFAP, CD34*, and *OLIG2* were used as cell-type specific marker genes for neurons, microglia, astrocytes, endothelial cells, and oligodendrocytes, respectively. ExonicRate: exonic mapping rate; RIN or RINcontinuous: RNA integrity number; AgeAtFirstADDxNum: age at first AD diagnosis; pmi: post-mortem interval; educ: education. (**c**,**d**) The variance partition analysis on microglial-specific gene expression residuals in MAYO (**c**) and ROSMAP (**d**) after de-convolution and covariate adjustment demonstrates that the microglial-specific residuals capture the microglial component (variance) and that the effects of other covariates and other cell types in the brain are removed.

**Figure S4.**
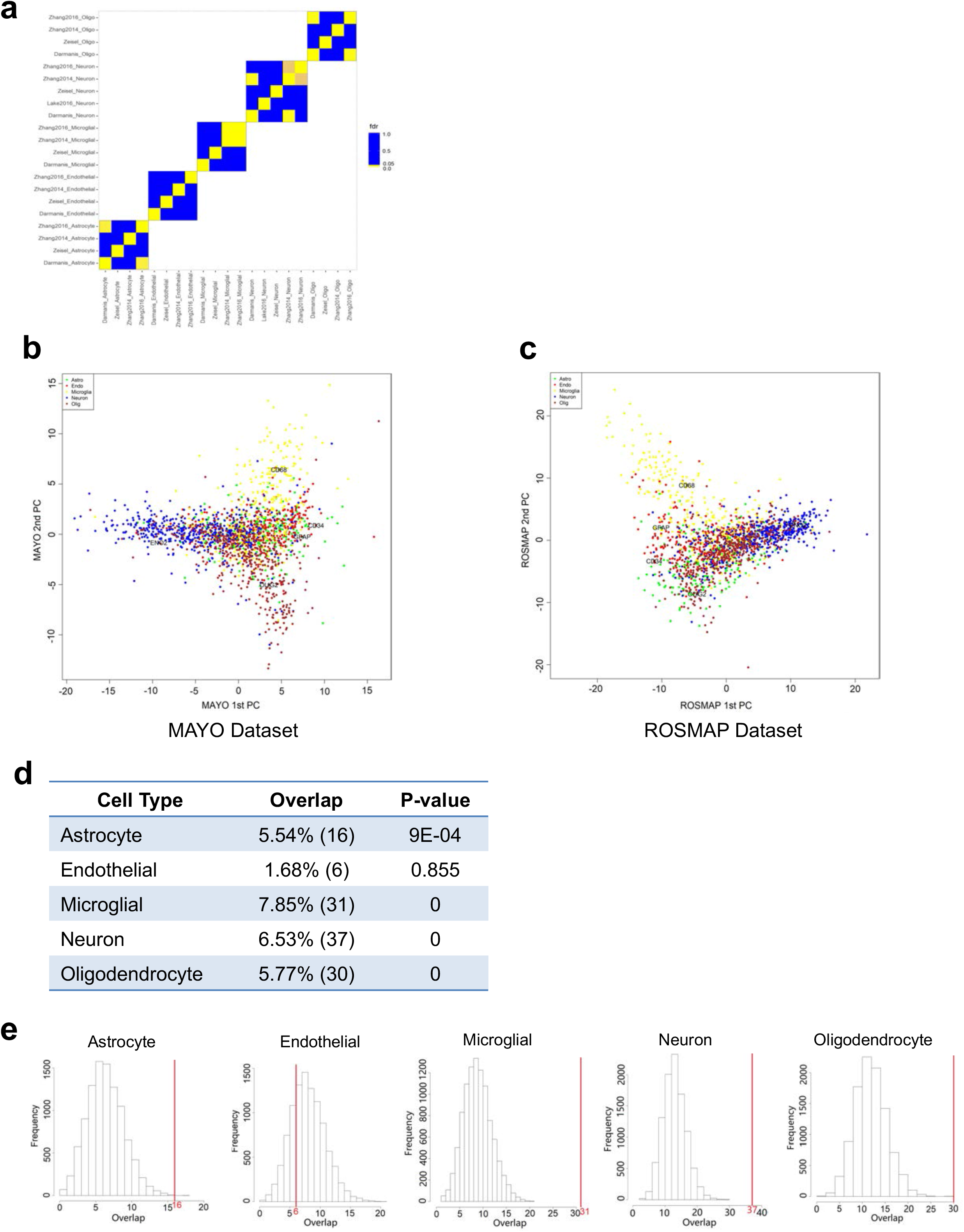
(**a**) Enrichment analysis of multiple multi-gene biomarker lists for the five main CNS cell types derived from various scRNAseq studies in control human brains shows no significant overlap among the studies. Significance was assessed by Fisher’s exact test with FDR cut-off of 0.05. (**b**,**c**) Principal component analysis (PCA) shows a prominent overlap of scRNAseq biomarker expression across the five main CNS cell types in AD residuals from the MAYO (**b**) and ROSMAP (**c**) datasets after covariate adjustment. (**d**) Enrichment analysis reveals significant overlap between scRNAseq biomarkers and AMP-AD Agora targets for neurons, microglia, astrocytes, and oligodendrocytes. The value in parentheses represents the number of genes overlapping between each biomarker list and the AMP-AD Agora targets. (**e**) Compared to randomly selected genes from the background overlapping with the AMP-AD Agora list, the number of overlapping genes between scRNAseq biomarkers and AMP-AD Agora targets (red vertical line along the distribution) is significantly higher for four cell types, including microglia. Significance was assessed by Fisher’s exact test with FDR cut-off of 0.05.

**Figure S5.**
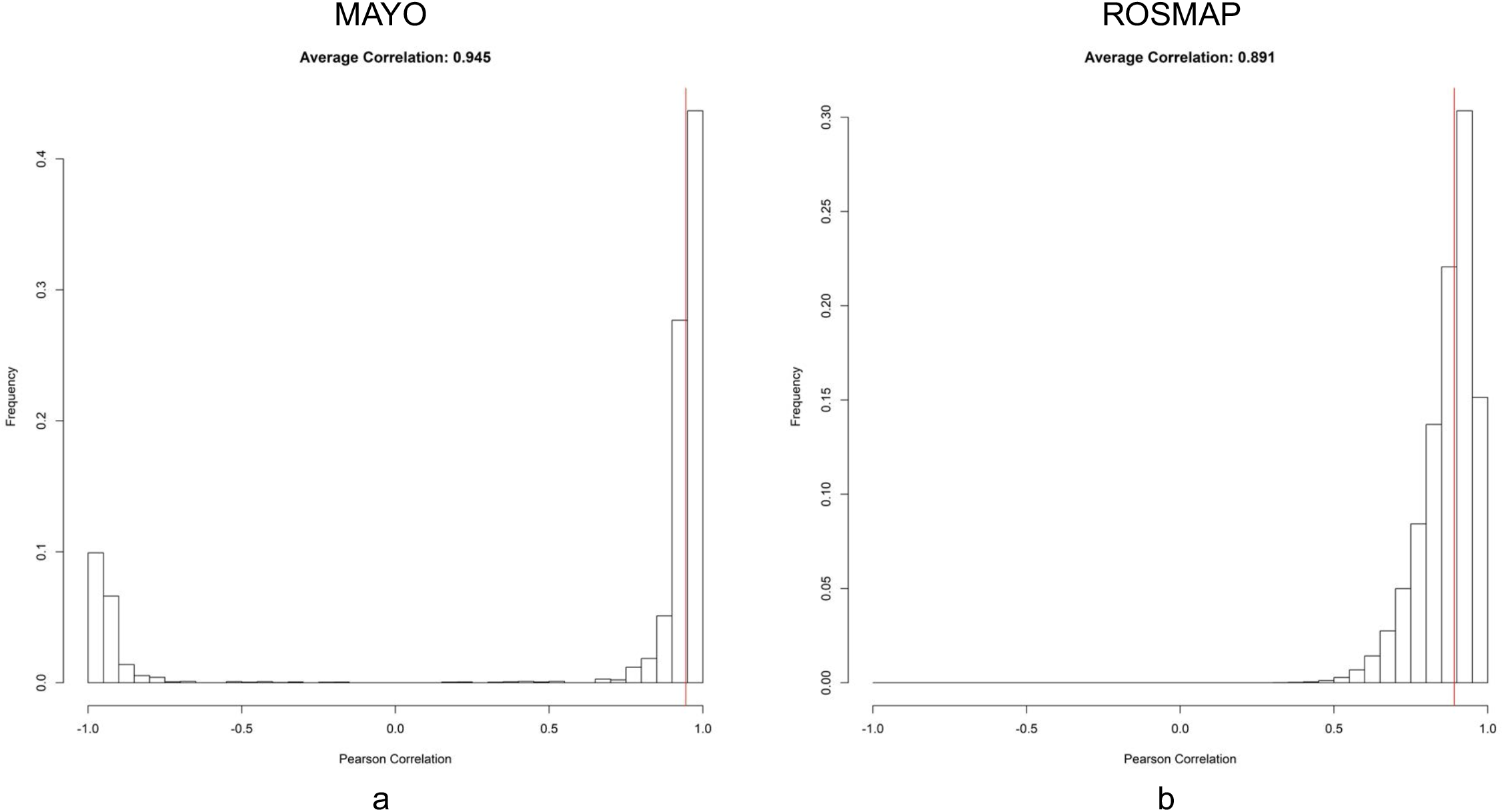
Distributions of Pearson correlation coefficients between our microglial-specific residual derived from *CD68* and the “pseudo” microglial-specific residuals derived from a randomly selected subset of microglial scRNAseq biomarkers by population-specific expression analysis (PSEA), for both the MAYO (**a**) and ROSMAP datasets (**b**).

**Figure S6.**
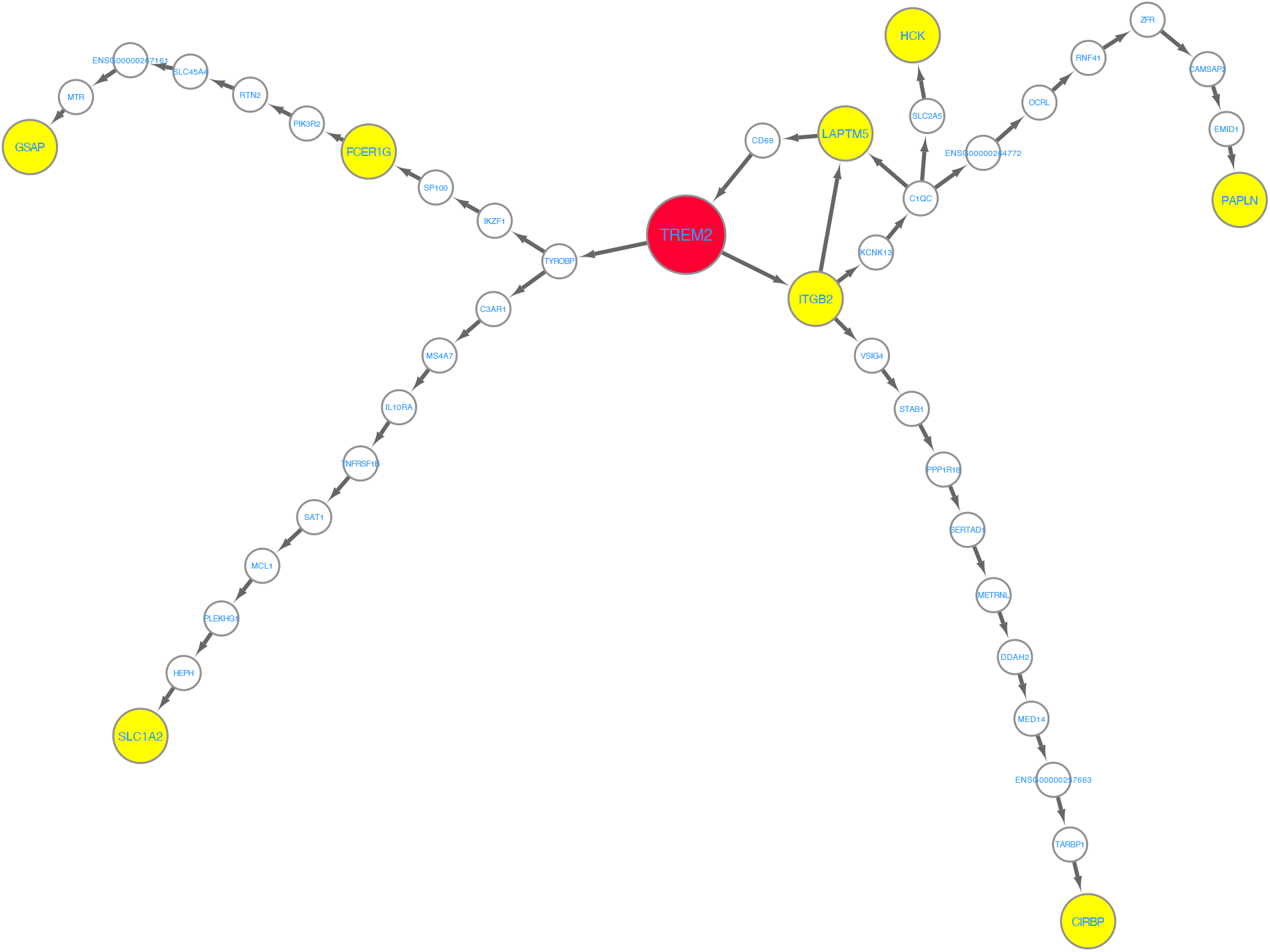
Regulatory pathway analysis reveals regulatory pathways between the validated targets of microglial phagocytosis in AD and *TREM2*. The shortest paths between each of the 9 targets and *TREM2* were extracted from each network and combine into a causal network.

**Figure S7.**
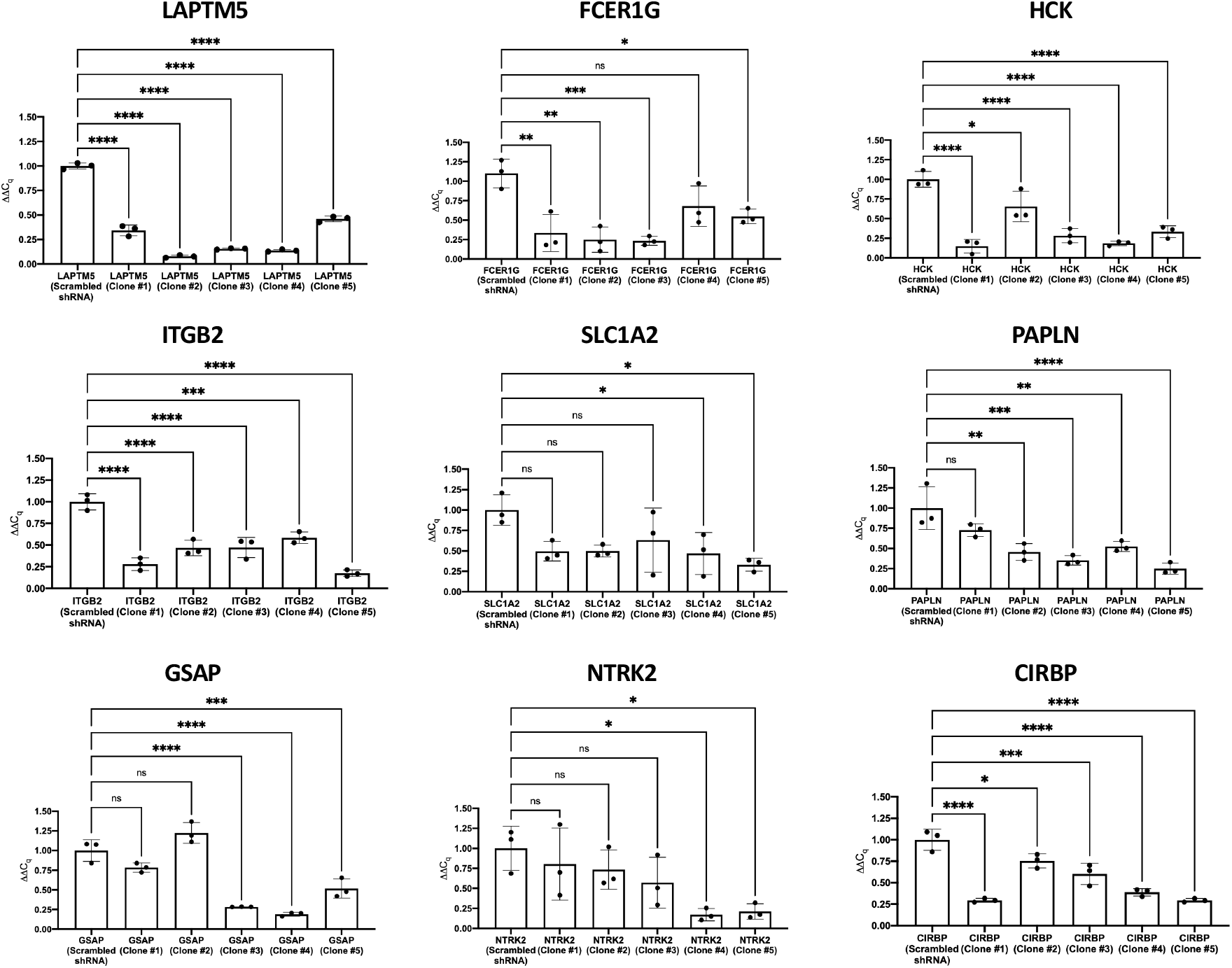
Knockdown of Key Drivers (KDs) by Lentiviral shRNA Clones. Using quantitative PCR (qPCR), mRNA expression for each of nine prioritized KDs was measured in HPMs following treatment with one of five targeting lentiviral shRNA clones. For each gene, a scrambled shRNA clone was used as a negative control. Data are expressed as mean ± SD of three experimental replicates (n = 3). Asterisks indicate data points that are statistically significant as compared to the negative control (* p < 0.05; ** p < 0.01; *** p < 0.001; **** p < 0.0001; ns = not significant).

**Table S1.** Microglial-specific differentially expressed (DE) gene expression signatures associated with AD in the MAYO and ROSMAP RNAseq datasets.

**Table S2.** Significantly enriched pathways associated with microglial-specific DE gene signatures in the MAYO and ROSMAP RNAseq datasets, indicating dysregulated microglial functions in AD. Pathway enrichment was assessed using Human ConsensusPathDB (CPDB).

**Table S3.** Enrichment analysis of microglial-specific DE gene signature against single-cell RNAseq (scRNAseq)-derived microglial DE gene signature.

**Table S4.** List of *cis*-eQTL genes associated with microglial component in the MAYO and ROSMAP datasets.

**Table S5.** Significantly enriched biological pathways associated with gene modules in the MAYO and ROSMAP microglial-specific co-expression networks.

**Table S6.** Significantly perturbed pathways associated with KD subnetworks of the 43 KDs for microglial phagocytosis in AD.

**Table S7.** Summary of statistical analyses of Aβ uptake by human primary microglial cells following shRNA knockdown of each of the 9 prioritized key driver targets.

**Table S8.** Knockdown percentage of KDs for the two best lentiviral shRNA clones.

## Acknowledgement

The authors thank the following funding resources which supported this study: NIH/NIA 1R56AG062620-01 and NIH/NINDS/Mayo Clinic U54NS110435 subaward to R.C.; NIH/NIA 1RF1AG057457-01 to R.C.; American Heart Association (AHA) Transformational Research Project Award (19TPA34910113) to P.T.R.; Ministry of Science and Technology (Taiwan) 109-2320-B-006-043-MY3 and 110-2320-B-006-021 to Y.M.K.

R.C. is the founder of INTelico Therapeutics LLC. R.C. and P.T.R. are co-founders of PATH Biotech LLC. This study is not supported by any funding from INTelico Therapeutics LLC or PATH Biotech LLC.

