## Supplementary material for "Novel Master Regulators of Microglial Phagocytosis and Repurposed FDA-approved Drug for Treatment of Alzheimer Disease": Online Method

**Materials and Methods**

### 1. AMP-AD Consortium Data Source

Data has been downloaded from the Accelerating Medicines Partnership – Alzheimer’s Disease (AMP-AD) consortium database hosted on the Synapse.org data portal (doi:10.7303/syn2580853).

#### 2. AMP-AD Mayo Clinic Cohort and Data Pre-Processing

**2.1) Mayo Clinic Transcriptome and Genome-Wide Genotype Data**

The Mayo Clinic (herein MAYO) transcriptome and genome-wide genotype datasets utilized in this study have previously been described[1-4]. The MAYO temporal cortex (TCX) RNA sequencing (RNAseq) data (Synapse ID: syn3163039) and genome-wide genotype data (Synapse ID: syn8650953) are available on the AMP-AD Knowledge Portal. We provide details on these datasets below.

**2.2) MAYO Cohort Participants**

Mayo Clinic RNAseq cohort includes RNAseq-based whole transcriptome data from TCX samples from 278 subjects with the following diagnoses: 84 AD, 84 progressive supranuclear palsy (PSP), 80 cognitive normal (CN) controls, and 30 pathologic aging. For this study, we utilized data from subjects with AD and CN samples. Subjects with AD had definite neuropathologic diagnosis according to the NINCDS-ADRDA criteria[5] and a Braak[6] neurofibrillary tangle (NFT) stage of ≥4.0. Control subjects each had Braak NFT stage of 3.0 or less, CERAD[7] neuritic and cortical plaque densities of 0 (none) or 1 (sparse) and each lacked any of the following pathologic diagnoses: AD, Parkinson’s disease (PD), dementia with Lewy bodies (DLB), vascular dementia (VaD), progressive supranuclear palsy (PSP), motor neuron disease (MND), corticobasal degeneration (CBD), Pick’s disease (PiD), Huntington’s disease (HD), frontotemporal lobar degeneration (FTLD), hippocampal sclerosis (HipScl), or dementia lacking distinctive histology (DLDH). Within the Mayo RNAseq cohort, all AD and PSP subjects were from the Mayo Clinic Brain Bank. Thirty-one control TCX samples were from the Mayo Clinic Brain Bank, and the remaining control tissue was from the Banner Sun Health Institute. All subjects were North American Caucasians. All disease subjects had ages at death ≥60 years; a more relaxed age cutoff of ≥50 years was applied for CN controls to achieve a sample size similar to that of the AD subjects, but we note there were only two additional control subjects with age at death below 60. A total of 76 control subjects and 79 AD subjects from the MAYO dataset were used in our analysis. We performed rigorous statistical testing to demonstrate that these samples are well balanced with respect to age at death (p-value=0.57) as well as sex (p-value=0.24) (Figure S2a,c). TCX samples underwent RNA extractions via the Trizol/chloroform/ethanol method, followed by DNase and Cleanup of RNA using Qiagen RNeasy Mini Kit and Qiagen RNase -Free DNase Set (Germantown, MD). The quantity and quality of all RNA samples were determined by the Agilent 2100 Bioanalyzer (Agilent Technologies, Santa Clara, CA). Samples had to have an RIN ≥5.0 for inclusion in the study. All of this work was approved by the Mayo Clinic Institutional Review Board. All human subjects or their next of kin provided informed consent.

**2.3) Mayo Clinic Genome-Wide Genotype Data**

Subjects in the Mayo Clinic RNAseq cohort underwent whole genome genotyping using the Illumina Infinium HumanOmni2.5-8 BeadChip, which delivers comprehensive coverage of both common and rare SNP content from the 1000 Genomes Project (minor allele frequency>2.5%) providing genotypes for 2,338,671 markers. The genotyping was done at the Mayo Clinic Medical Genome Facility. Whole genome genotype calls were made using the auto-calling algorithm in Illumina’s BeadStudio 2.0 software, subsequent to which they were converted into PLINK formats for analysis[8].

**2.4) MAYO Genotype Data Quality Control**

Genome-wide genotypes were obtained for all subjects in the MAYO RNAseq study using Illumina Omni 2.5 Beadchips. Samples were checked for discordant sex, and we identified and excluded 2 AD subjects for this reason. Subjects were assessed for heterozygosity rates >3 SD from the mean. One AD sample had high heterozygosity with respect to the mean, indicating possible sample contamination, and 3 samples (2 controls and 1 AD) had low heterozygosity with respect to the mean, indicating either divergent ancestry or consanguinity; these 4 samples were also excluded from the analysis. The dataset was then filtered to include only autosomal SNPs. PLINK was used to identify any sample duplicates or related pairs of subjects. Two pairs of samples were identified as >3^rd^ degree relatives; for each pair, the sample with the lower SNP call rate was excluded. The dataset was further filtered to remove complex genomic regions (chr8:1-12,700,000; chr2:129,900,001-136,800,000; chr17:40,900,001-44,900,000; and chr6:32,100,001-33,500,000) and linkage disequilibrium (LD) pruned using the SNPRelate (v1.4.2) package in R (v3.2.3) [9], implementing an LD threshold of 0.15 and a sliding window of 1E-07 bp. Remaining SNPs and subjects were analyzed using EIGENSOFT[10] for population outliers. Two samples were identified as population outliers using the default parameter of >6 SD from the mean on any of the top 10 inferred axes following 5 iterations, and they were removed from further analysis.

**2.5) Mayo Clinic RNAseq Data**

Mayo Clinic RNAseq samples were randomized across flowcells, taking into account age at death, sex, RIN, Braak stage and diagnosis. Library preparation and sequencing of the samples were conducted at the Mayo Clinic Medical Genome Facility Gene Expression and Sequencing Cores, as previously described[11]. The TruSeq RNA Sample Prep Kit (Illumina, San Diego, CA) was used for library preparation from all samples. The library concentration and size distribution were determined using an Agilent Bioanalyzer DNA 1000 chip (Agilent Technologies). Three samples were run per flowcell lane using barcoding. All samples underwent 101 base-pair (bp), paired- end sequencing on Illumina HiSeq2000 instruments. Base-calling was performed using Illumina’s RTA 1.17.21.3. FASTQ sequence reads were aligned to the human reference genome using TopHat 2.0.12[12] and Bowtie 1.1.0[13], Subread 1.4.4 was used for gene counting[14]. FastQC[15] was used for quality control (QC) of raw sequence reads, and RSeQC[16] was used for QC of mapped reads.

**2.6) MAYO RNAseq Data Quality Control**

All MAYO RNAseq samples had percentage of mapped reads ≥85%. In the first step, we used R statistical software (R Foundation for Statistical Computing, version 3.2.3) to transform the raw read counts to counts per million (CPM), which were then log2-normalized. Mean expression levels of Y chromosome genes with non-zero counts were plotted to identify any samples with deviation from expected expression based on recorded sex; the same 2 AD samples as mentioned in the genome-wide genotype QC (section 2.4) were identified with discordant sex and excluded. In the second step, raw read counts were normalized using Conditional Quantile Normalization (CQN) via the Bioconductor package[17], accounting for sequencing depth, gene length, and GC content. Before running CQN normalization, GC content was calculated via Repitools in the Bioconductor package[18] and sequencing depth was calculated as the sum of reads mapped to genes. Genes with non-zero counts across all samples were retained and principal component analysis (PCA) was performed using the prcomp function implemented with R statistical software. Principal components 1 and 2 were plotted and no outliers (>6 SD from mean) were identified.

#### 3. AMP-AD ROSMAP Cohort and Data Pre-Processing

**3.1) ROSMAP Genome-Wide Genotype and Transcriptome Data**

The Religious Orders Study and Memory and Aging Project (herein ROSMAP) dataset dorsolateral prefrontal cortex (DLPFC) gene expression (RNAseq BAM files), genotypes, and clinical covariates were downloaded from the Synapse.org data portal (Synapse IDs for the respective data types: syn4164376, syn3157325, and syn3191087) using the synapseClient R library[19]. Requests for ROSMAP data can be made at https://www.radc.rush.edu/.

**3.2) ROSMAP Cohort Participants**

The ROSMAP dataset contains two cohorts: the Religious Orders Study (ROS) and the Memory and Aging Project (MAP)[20]. Both ROS and MAP are longitudinal clinical-pathologic cohort studies of aging and dementia run by the Rush Alzheimer’s Disease Center in Chicago, IL. In both studies, all participants enroll without known dementia and agree to annual clinical evaluation and brain donation as a condition of entry. ROS has enrolled individuals from religious orders from across the United States starting in 1994, and MAP has enrolled lay persons from across northeastern Illinois since 1997. Each study annually administers a battery of 21 cognitive performance tests, 19 of which are in common. Alzheimer's disease status is determined by a computer algorithm based on cognitive test performance with a series of discrete clinical judgments made by both a neuropsychologist and a clinician. First, subjects are categorized as not cognitively impaired (NCI, if diagnosed without dementia), mild cognitive impairment (MCI), or Alzheimer’s disease (AD). Diagnoses of dementia and AD conform to standard definitions[20]. Next, a clinician reviews all cases determined by the algorithm to render a diagnosis blinded to data collected in prior years. In addition to dementia, 5 other diagnoses are determined by this approach, including stroke, cognitive impairment due to stroke, parkinsonism, Parkinson's disease, and depression. Most of these other diagnoses are determined by self-report. Upon death, a summary diagnosis is made by a neuropsychologist blinded to post-mortem assessment. The post-mortem neuropathologic evaluation performed includes a uniform structured assessment of AD pathology, cerebral infarcts, Lewy body disease, and the other pathologies common in aging and dementia (e.g., vascular dementia or frontotemporal dementia). The evaluation procedures follow those outlined by the pathologic dataset recommended by the National Alzheimer’s Disease Coordinating Center. Pathologic diagnoses of AD use NIA-Reagan and modified CERAD criteria[21], and the evaluation of neurofibrillary pathology uses Braak staging[22]. The ROS and MAP studies are both conducted by the same clinical and pathologic data collection teams, with extensive item-level harmonization allowing the data to be efficiently merged. A total of 194 CN subjects and 212 AD subjects from the ROSMAP dataset were used in our analysis. Regarding the ROSMAP cohort, it has been noted that the range of age at death is broad but restricted to the older segment of the age distribution of the North American and that age and sex are important confounders when performing any analyses of ROS and MAP data[23]. We observed this variance in the age at death but found no significant difference in sex among the ROSMAP subjects used in our analysis (Figure S2b,d). To address this imbalanced age distribution, we performed covariate adjustment (Section 5) and confirmed of the effects of this confound by variance partition analysis (VPA) before and after covariate adjustment (Figure S3b,d).

**3.3) ROSMAP Genotype Data Processing**

PLINK 2.0[24] was used to perform operations on the genotype files, and positions were converted from hg18 to hg19 (http://genome.ucsc.edu/cgi-bin/hgLiftOver). Picard[25] was used to sort the resulting genotype files, and samples were removed using PLINK 2.0 if they had variants with >2% missing values, minor allele frequency <1%, Hardy-Weinberg equilibrium <10E-6, or inbreeding coefficient >0.15. We started with 750,173 variants in 1,708 individuals; after quality control, 736,073 variants in 1,091 individuals remained.

### 3.4) ROSMAP RNAseq Data Processing

The RNA-seq BAM files were sorted using samtools[26] and converted to FASTQ files using the SamToFastq function[25]. RAPiD[27] was used to generate a count matrix for the gene expression data and a vcf file for each sample aligned to hg19 from the FASTQ files. ROSMAP RNAseq read count expression data was normalized using log2 counts per million (CPM) and the TMM method[28] implemented in edgeR[29]. Genes with over 1 CPM in at least 30% of the experiments were retained. We then used precision weights as implemented in the voom function from the limma[30] R package to further normalize the gene counts.

**4. MAYO and ROSMAP Genotype Data Imputation**

After quality control of both datasets, we used 1000 Genomes Project[31] data and IMPUTEv2[32] to impute untyped variants. Imputed variants were removed if they failed any of the previously listed quality control criteria or had information scores <0.6. After imputation, we had 7,132,687 variants in MAYO and 9,333,139 variants in ROSMAP.

### 5. Deconvolution of RNAseq Data into Microglial-Specific Expression Residuals

Central nerve system (CNS) tissue consists of various cell types, including neurons, endothelial and glial cells. To discover key drivers specific to a single cell type in the CNS and study their contribution to AD in that specific cell type, we utilized validated single-cell marker genes to directly deconvolve bulk-tissue gene expression data into cell type-specific gene expression for five primary cell types in the CNS: neurons, microglia, astrocytes, endothelia, and oligodendrocytes. In this study, we focused on investigating the role of microglial cells in AD as recent genome-wide association studies linked AD risk loci with microglial functions[33-36], which defined the overall cellular response to the accumulation of Aβ[37, 38]. After normalizing the bulk-tissue RNAseq data but before covariate adjustment, we performed variance partition analysis (VPA) [39] to evaluate the contributions of cell-type specific markers as well as demographic, clinical, and technical covariates (such as batch effects) to the gene expression variance before performing any covariate adjustment in MAYO and ROSMAP cohorts(Figure S3a,b). The cell-type specific marker genes used for neurons, microglia, astrocytes, endothelial cells, and oligodendrocytes were *ENO2*, *CD68*, *GFAP, CD34*, and *OLIG2,* respectively, as previously published [40]. The VPA results reflect the prominent effect of CNS cell types on the variance of the brain RNAseq data. In the MAYO dataset, the additional covariates used in the VPA included exonic mapping rate calculated by RNAseQC[41], RNA integrity number (RIN), sequencing batch, diagnosis, age at death, tissue source, *APOE* genotype, and sex. In the ROSMAP dataset, we were able to include the same covariates with the exception of tissue source and the addition of age at first AD diagnosis, post-mortem interval (PMI), education, and study (ROS or MAP). Next, we performed covariate adjustment and deconvolution using the PSEA method[42] in each dataset, calculating gene expression residuals using a linear regression model to adjust the normalized bulk-tissue expression data with demographical and technical covariates as well as the cell-type specific markers. Cell-type specific gene expression, including the microglial-specific component, was directly derived by adding the estimated variance of each cell type to the residual, avoiding the need to first estimate the cell population from bulk tissue data, which could induce approximation errors. After covariate adjustment, we then repeated VPA in the microglial-specific residuals of each dataset to demonstrate that our deconvolution and covariate adjustment methods properly capture the microglial component while removing potential confounds such as batch effect, age, and sex(Figure S3c,d). Finally, to justify the use of single cell-type specific markers for deconvolution by the PSEA method, we performed a set of analyses comparing multiple cell-type specific biomarker lists (derived from existing scRNAseq studies) to each other, to our AD residuals, and to the AMP-AD Agora list of potential therapeutic targets in AD (Online Method 5.2), as well as a robustness analysis demonstrating that our microglial-specific residual derived from *CD68* expression represents a robust microglial component in the bulk-tissue RNAseq data when compared to random selections of multi-gene microglial biomarkers derived from these scRNAseq datasets in AD. The final microglial-specific expression residual data available for further analysis included 19,885 genes from 76/79 CN/AD samples in MAYO and 20,276 genes from and 194/212 CN/AD samples in ROSMAP, with 18,408 genes common to both datasets which are comparable with processed residuals of the same cohorts on AMP-AD knowledge portal[43].

### 5.1) Deconvoluted Microglial-Specific Gene Expression vs scRNAseq

Although single-cell RNA sequencing (scRNAseq) data has significantly advanced our understanding of cellular heterogeneity [44-46] and de-novo discovery of cell populations [47, 48], as well as spurred developments of various computational analysis tools [49], the performance of network inference and key driver discovery from scRNAseq data is still very poor, yielding a significant amount of uncertainty in inferred network models [50, 51]. This limitation of scRNAseq data in network inference is mainly attributed to its high amount of missing gene expression measures (dropouts) and the immaturity of the network methods dealing with such enormous amount of missing data. On the other hand, various methods of deconvolution of bulk-tissue RNAseq data into single cell type gene expression have become increasingly popular in recent years as a complementary solution to the missing values in scRNAseq data [52-64]. The core assumption of these deconvolution methods is that gene expression in bulk-tissue data is equivalent to the averaged gene expression of each cell type weighted by its relative population in the tissue. Thus, these methods decompose bulk-tissue RNAseq data into gene expression of individual cell types by estimating the relative cell populations in the tissue via cell type-specific biomarker genes. After deconvolution, the variance of the deconvolved gene expression of each cell type becomes orthogonal to other cell types and can be analyzed independently[42].

**5.2) Rationalization and Validation of Single-Gene Biomarkers for Bulk-Tissue RNAseq Deconvolution**

The rational of using single-gene biomarkers in the above over multi-gene biomarkers derived from scRNA-seq data is as follows: 1) multi-gene biomarkers derived from various scRNA-seq studies in human brain under control conditions [65-69] shows non-significant (FDR>0.05, Online Methods) overlap (Figure S4a, 1/0/1/1 significant pair out of 6 study-pairs in Asctrocyte/Endothelial/Microglial/Oligodendrocyte types and 2 significant pairs out of 10 study-pairs in Neuron), indicating lack of robustness and consensus in these biomarkers derived from these studies; 2) significant overlap of scRNA-seq-derived biomarkers expression in ROSMAP and MAYO AD samples by PCA analysis (Figure S4b,c), indicating that the majority of scRNAseq-derived biomarker gene expression is convoluted and reflecting potential interactions between different cell types under the AD condition; 3) Significant overlap between scRNA-seq-derived biomarkers and AD therapeutic targets in the AMP-AD AGORA knowledge portal (https://agora.ampadportal.org/genes) (Figure S4d,e). This overlap rate is significantly higher than randomly selected genes from the background overlapping with the AGORA list, indicating that scRNA-seq-derived biomarkers may play a significant role in AD pathology. For these reasons, multi-gene biomarkers of cell type are not ideal for adjusting the bulk-tissue gene expression variance by population-specific expression analysis (PSEA). By contrast, our single-gene biomarkers are derived from biological knowledge and validated by others [40]. Moreover, our single-gene biomarker had no overlap with AD therapeutic targets in AGORA, which make them good candidates for PSEA. 4) Further, our single-gene biomarker derived microglial-specific residual is significantly (p-value<2.2E-16, Figure S5a,b) correlated with the “pseudo” microglial-specific residuals derived from randomly selected subset of scRNA-seq biomarkers[65-69] by PSEA, indicating that our single-gene biomarker derived microglial-specific residual represents a robust microglial component in the bulk-tissue RNAseq data for microglial-specific therapeutic target discovery in LOAD.

**5.3) Validation of microglial-specific DE signatures derived from the Bulk-Tissue RNAseq Deconvolution vs scRNA-seq data in AD**

To further demonstrate that our microglial-specific DE signatures derived from deconvoluted microglial gene expression significantly correlated with scRNAseq-derived microglial signatures in AD, we employed the sampling-based method [63] to calculate enrichment of ROSMAP-microglial and MAYO-microglial DE genes with 21 independent scRNAseq-derived microglial signatures[31, 63-72]. We found 3 and 20 scRNAseq-derived microglial signatures were enriched by our MAYO- and ROSMAP-microglial DE signature respectively (Table S3). Interestingly, both MAYO- and ROSMAP-microglial DE signatures are significantly (p-value=0.021 and 0.030) enriched for the scRNAseq-derived DE signature of disease-associated microglia (DAM)[72]. Recent evidence suggests that DAM may play a protective role by detecting neurodegeneration-associated damage within the CNS mediated by TREM2 signaling pathway[71]. In addition, MAYO-microglial DE signature are enriched for microglial response to injury[70]. ROSMAP-microglial DE genes are enriched for signatures associated with axon tract-associated microglia (ATM)[70], aged brain[70], LPS-induced activation[69], increased neuroinflammation[68], cell culturing[67], and neurological disorders including ALS[66], glioma[31, 65], epilepsy[31], and amyloid plaques in AD[63, 64]. In summary, our deconvoluted microglial gene expression captured the activated microglial state associated with AD pathology.

### 6. Computational Analysis of Microglial-Specific Gene Expression Data

### 6.1) eQTL Analysis

Expression quantitative trait locus (eQTL) analysis was performed using the R package MatrixEQTL v2.1.1[70] using QCed genotypes and normalized and covariate-adjusted cell-type specific expression residuals. *cis*-eQTL analysis considered markers within 1 Mb of the transcription start site of each gene. False discovery rates (FDR) were computed using the Benjamini–Hochberg procedure [71].

### 6.2) Differential Expression (DE) Analysis

Using linear models, as implemented in the limma R package [30], we interrogated the cell-type specific residual expression data for genes differentially expressed between AD cases and healthy controls. Significance was assessed using FDR<0.05.

**6.3) Co-Expression Network Analysis**

Co-expression networks were constructed using the coexpp R package[72]. A soft thresholding parameter value of 6.5 was used to power the expression correlations.

**6.4) Co-Expression Module Selection**

We identified 6 modules (M1, M8, M11, M12, M33, M43) were significantly enriched for microglial-DE signature and two modules (M6 and M45) significantly enriched for microglial cell markers in MAYO dataset (FDR<0.05) and 11 modules (M3, M4, M6, M7, M12, M23, M30, M35, M37, M38 and M40) were significantly enriched for microglial-DE signature and two modules (M8 and M11) enriched for microglial cell markers in ROSMAP.

We also noted that microglial DE-enriched modules in ROSMAP were also significantly enriched for markers of other cell types, indicating microglial-involved cell-cell interactions were captured by our co-expression network. It is well established that microglia interact with other brain cells in modulating amyloid pathology and neuroinflammation in mouse models of Alzheimer’s disease[68]. Failing to account for these interactions will result in a compromised network model, however, over-counting these interactions will decrease the network specificity to microglial cells. Therefore, we only considered microglial-interacting cell type(s) whose marker genes are the most significantly enriched (FDR<10e-4) by the same modules that are also the most significantly enriched (FDR<10e-4) for microglial-DE signature (Figure 3a). Across the two co-expression networks, we noted that only two modules (M6 and M23) in ROSMAP were enriched for both microglial-DE signature and astrocyte markers with FDR<10e-4, suggesting a strong signal of microglial-astrocyte interactions in our microglial-specific gene expression component. Consequently, in addition to the modules enriched for the microglial-DE and microglial markers, we added two modules (M4 and M39) and two modules (M6 and M23) enriched for astrocyte cell marker genes in MAYO and ROSMAP respectively. Genes in these additional modules reflecting astrocyte-microglia interactions accounted for only 13.9% (583 genes) and 16.9% (873 genes) of the MAYO- and ROSMAP-microglial co-expression network respectively. To form final seeding set of genes for input into the causal predictive network modeling in each cohort respectively, we selected 10 modules (6 enriched for microglial-DE, 2 enriched for microglial cell markers, and 2 enriched for astrocyte cell markers) in MAYO-microglial co-expression network and 13 modules (11 enriched for microglial-DE, including the 2 also enriched for astrocyte cell markers and 2 enriched for microglial cell markers) in the ROSMAP-microglial co-expression network.

**6.5) Key Driver Analysis**

To perform Key Driver Analysis (KDA), we used the KDA R package [73] (KDA R package version 0.1). The package first defines a background sub-network by looking for a neighborhood k-step away from each node in the target gene list in the network. Then, stemming from each node in this sub-network, it assesses the enrichment in its k-step (k varies from 1 to K) downstream neighborhood for the target gene list. In this analysis, we used K = 6.

**6.6) Key Driver Prioritization**

To prioritize key drivers, we first calculated the following three metrics for every key driver:

1) Robustness_Score: We calculated how many datasets, i.e., MAYO and ROSMAP cohorts, by which a key driver was replicated.
2) TargetType_Score: We calculated how many categories of target source predicted a gene as KD in KDA. In this study, we used 3 types of target source in KDA: DE genes, co-expression modules and overlapping genes of DE gene with modules.
3) TargetSet_Score: We calculated the total number of different target sources predicted a gene as a KD in KDA.

Next, we prioritized all key drivers according to three measurements. We firstly ranked the key drivers according to *Robustness_Score* in descending order, then ranked by *TargetType_Score* in descending order, and lastly ranked by *TargetSet_Score* in descending order.

### 6.7) Predictive Networks Reconstruction

Our causal network pipeline starts by integrating genome-wide genotyping and RNAseq data generated from the dorsolateral prefrontal cortex of 612 subjects in ROSMAP[23, 74-76] and from the temporal cortex of 266 subjects in MAYO [40, 77, 78] in the Accelerating Medicines Partnership - Alzheimer's Disease (AMP-AD) consortium, spanning the complete spectrum of AD clinical and neuropathological traits. We processed matched genotype and RNAseq data separately in each cohort.

Although the co-expression network modules capture highly co-regulated genes operating in coherent biological pathways, these modules do not reflect the probabilistic causal information needed to identify key driver genes. Bayesian networks (BNs)[79] are a long-standing form of statistical network modeling used to reverse-engineer probabilistic causality among variables; with the development of high-throughput sequencing technology, BNs have been widely used to infer causal gene regulatory networks in different diseases [80-85]. Recent studies have applied BNs to infer molecular mechanisms and key drivers in Alzheimer’s disease[86, 87].

However, BNs have significant limitations with respect to inferring opposite causality given the symmetry of joint probability. Recent work has demonstrated that bottom-up causality inference can accurately distinguish true causality from opposite causality in equivalent classes[88]. In this study, we developed a novel computational network model, called predictive network modeling, by integrating conventional (top-down) Bayesian networks with bottom-up causality inference to address the problem of opposite causality inference in BN modeling. Here, our causal predictive network pipeline incorporates multi-scale omics data, including genotypes and transcriptomic profiles, in the MAYO and ROSMAP datasets (deconvoluted microglial-specific residuals) to build causal predictive networks separately in both datasets.

**6.8)** **CPDB Pathway Enrichment Analysis**

We downloaded pathways from ConsensusPathDataBase (CPDB)[89]. Given a set of genes, we performed enrichment analysis of each pathway over this set of genes by Fisher’s exact test.

**6.9) Genetic Regulatory Pathways among 9 KDs and TREM2**

The shortest directed paths between the 9 validated key drivers and TREM2 are derived from MAYO- and ROSMAP-microglial predictive networks and combined into a network elucidating their regulatory relationship.

**7. Network-driven Inverse Gene Expression Score for Drug Repurpose**

We developed a network-driven inverse gene expression score for drug repurposing. We first derived the differential expression signature (|z-score|>2) from the total 720,216 drug-based perturbation profiles in level-5 CMAP/LINCS database over 230 cell lines derived from 21 tissue types (including unknown and central nervous system) and computed geometric similarity and rank-sum test statistics between CMAP perturbation signature (vector of z-scores) and ROSMAP- and MAYO-microglial DE signature (vector of log-Fold change values) for the genes in the downstream subnetwork of every 9 validated target in ROSMAP and MAYO networks respectively. A repurposed perturbation is considered a ‘hit’ against a target gene if the geometric similarity is significantly (FDR<0.05) negative in the rank-sum test, indicating this perturbation significantly reversed the gene expression of the downstream pathways of a target gene from AD to CN. As a result, we identified a total of 26,375 and 59,519 significant perturbation hits (out of total 720,216 perturbations in CMAP) in MAYO and ROSMAP cohorts, respectively, with 11,650 overlapped perturbations over 207 cell lines. Out of these perturbations, 1425 and 762 significant (FDR < 0.05) perturbation hits are derived from six CNS cell lines in CMAP for MAYO and ROSMAP cohorts, respectively, with 292 hits overlapped. These 292 overlapped perturbations are mapped to 228 (out of total 33,609) drugs in CMAP, indicating their robust reversal effects on downstream gene expression of *SLC1A2* in ROSMAP- and MAYO-microglial network.

**8. Human primary microglial (HPM) cell culture**

Passage two HPM cells (Lot # 1614453-01) from an unaffected (i.e., non-AD) Caucasian 32-year-old female subject were obtained from Celprogen, Inc. (Torrance, CA, USA). The cells were expanded up to passage five in T175 flasks (Sarstedt, Inc., [Nümbrecht, Germany](https://www.google.com/search?client=safari&rls=en&q=N%C3%BCmbrecht&stick=H4sIAAAAAAAAAOPgE-LVT9c3NEypzE1Oiq8qV-LUz9U3ME43qTTUMsoot9JPzs_JSU0uyczP088vSk_My6xKBHGKrTJSE1MKSxOLSlKLihVy8pPBwotYufwO78lNKkpNzijZwcoIACX8bpBgAAAA&sa=X&ved=2ahUKEwiLx-bJ4tHxAhVIpZ4KHVMGAI4QmxMoATAVegQIERAD)) with ventilated caps and were maintained at 37 °C in a humidified 5% CO_2_ environment. HPMs were grown in Human Microglia Primary Tissue Culture Complete Growth Medium containing antibiotics (Celprogen, CAT#M37089-01) and supplemented with 10% Fetal Bovine Serum (Invitrogen, Waltham, MA, USA). Culture medium was replaced with fresh medium every 2 to 3 days. Cells were subcultured with 0.25% trypsin-EDTA upon reaching 95% confluence. All cell-based experiments to assess microglial phagocytosis were conducted on HPMs at passage 6.

**9. Knockdown of the nine key drivers in human primary microglia (HPM) using lentivirus-mediated RNA interference (RNAi)**

Genetic knockdown of target genes was performed using lentiviral clones that produced targeting shRNAs. To select viral clones that reduced expression of specific gene targets by approximately 70%, we tested knockdown efficiency of five unique clones per target gene (Sigma-Aldrich, St. Louis, MO, USA). A non-targeting shRNA producing lentivirus clone (MISSION pLKO.1-puro; Sigma-Aldrich, St. Louis, MO) was used against each specific target as a negative control. On day one, we plated 16,000 HPM cells per well with three wells per each lentiviral clone in flat-bottomed 96-well plates (Greiner Bio-One, Monroe, NC, USA; item # 655090). The following day, the cells were incubated with the lentiviral clones with a multiplicity of infection (MOI) of one in fresh cell culture medium supplemented with polybrene (Hexadimethrine bromide; Sigma-Aldrich, St. Louis, MO, USA) to a final concentration of 8 µg/mL. Sixteen hours later, cell culture medium containing viral particles was aspirated and fresh medium was added to each well. The next day, the regular cell culture medium was replaced with selection medium consisting of 2 µg/mL puromycin (Thermo Fisher Scientific, Waltham, MA, USA). The transfected cells received a fresh selection medium every second day for a period of two weeks.

At the end of the two-week transfection period, RNA was extracted from each well using an Aurum Total RNA Mini Kit (Bio-Rad Laboratories, Hercules, CA, USA) according to manufacturer’s instructions. The extracted RNA was then used to synthesize cDNA using a High-Capacity cDNA Reverse Transcription Kit with RNase Inhibitor (Thermo Fisher Scientific, Waltham, MA, USA). Quantitative PCR (qPCR) was performed to quantify the relative expression of each target using TaqMan Fast Advanced Master Mix (Thermo Fisher Scientific, Waltham, MA, USA) using a CFX96 Real-Time System thermocycler (Bio-Rad Laboratories, Hercules, CA, USA). The expression levels of *HPRT1* gene (i.e., the housekeeping gene) were used to calculate relative gene expression for each target via the ΔΔCq method. Based on these data (Supplementary Figure S7), the two lentiviral clones that had the highest efficiency of target gene expression knockdown (Supplementary Table S8) were selected for downstream analyses.

**10. Aβ1-42 uptake assay**

**10.1) Aβ1-42 uptake assay in Key Driver Knockdown in HPM**

We measured uptake of fluorescent Aβ1-42 in HPM cells transfected with lentiviral clones that produced targeting shRNA constructs. Transfected HPMs were cultured for three weeks to achieve sufficient downregulation of target gene expression. At this time, culture medium was aspirated and fresh growth medium containing HiLyte Fluor 488-labeled human Beta-Amyloid 1-42 (1.5 µg/mL; AnaSpec, Fremont, CA, USA) was added. Cells were then incubated for 2 h at 37°C in a humidified environment. HPMs were then washed three times with 1X PBS, fixed in 4% paraformaldehyde (PFA) for 15 min, and permeabilized using 0.01% Triton X-100 for 5 min. Microglial nuclei and cytoskeleton were labeled using Hoechst 33342 (1:1000 dilution in PBS, pH 7.4) and phalloidin (1:1000 dilution in PBS, pH 7.4), respectively. The imaging of stained cells was performed using an Operetta CLS High-Content Analysis System (PerkinElmer, Inc., Waltham, MA, USA). To obtain images for analysis, the following settings were used: Two-peak autofocus, 20x Air, NA 0.4 objective, Hoechst 33342 (2ms, power at 50%, 2µm height), Alexa 647 (50ms, power at 50%, 2µm height), Alexa 488 (1200ms, power at 50%, 2µm height). The mean fluorescence intensity per cell per well (MFCMFI), a parameter that reflects the quantity of fluorescently labeled Aβ1-42 internalized by microglia (i.e., phagocytosis), was measured using Harmony 4.5 software (PerkinElmer, Inc., Waltham, MA, USA). The two-sided, un-paired Wilcoxon rank-sum test was used to determine statistical significance.

**10.2) Aβ1-42 uptake assay in HPM treated with Riluzole**

In this experiment, primary cultures of human microglia were challenged with 1uM of HiLyte Fluor 488-labeled human Beta-Amyloid 1-42 (AnaSpec, Fremont, CA, USA) for 24 h. At 24 h, riluzole at 10uM was added to the culture media containing Aβ1-42 and intracellular fluorescence associated with Aβ1-42 was measured 24 h later. Aβ1-42 uptake in primary cultures of human microglia was measured by removal of medium containing fluorescent Aβ1-42 and addition of 200 ul of ice-cold PBS to stop cellular uptake mechanisms. At this time, cells were solubilized with 100 ul of 0.2% sodium dodecyl sulfate for 10 minutes. The human Aβ1-42 signal was measured using a fluorescent assay plate reader at an excitation wavelength of 450 nm and an emission wavelength of 535 nm. All samples were corrected for background fluorescence.

**11. Western Blot Analysis**

Western blotting on HPMs was performed according to a previously described method [90]. Briefly, whole cell lysates from primary cultures of human microglia were prepared by exposing cells to 2.0 ml of modified radioimmunoprecipitation (RIPA) buffer (50 mM Tris-HCl, pH 7.4, 150 mM NaCl, 1.0 mM EGTA, 1.0 mM sodium-*o*-vanadate, 1% (v/v) Nonident P-40, 0.25% (m/v) sodium deoxycholate, 0.1% sodium dodecyl sulfate, 200 μM phenylmethylsulfonyl fluoride, and 0.1% protease inhibitor cocktail (Millipore-Sigma)). Cells were then gently rocked for 15 min at 4°C to permit lysis. Cell suspensions were collected and centrifuged at 3000 x *g* for 15 min at 4°C to remove cellular debris. The supernatant was then collected and centrifuged at 100,000 x *g* for 1 h at 4°C. The resultant pellet was resuspended in 10 mM Tris buffer, pH 8.8, and stored at -20°C until further use. Protein concentration of the cell lysates was determined via Bradford’s protein assay. For western blotting, cell lysates were heated at 37°C for 30 min under reducing conditions (i.e., 2.5% (v/v) 2-mercaptoethanol (Millipore-Sigma)) in 1X Laemmli sample buffer (Bio-Rad, Hercules, CA) for SLC1A2 detection. All cell lysate samples were resolved on a 4-12% Criterion™ XT Bis-Tris polyacrylamide gel (Bio-Rad). Following SDS-PAGE and transfer, PVDF membranes blocked for 1 h in Superblock™ TBS blocking buffer (ThermoFisher) and then incubated overnight at 4°C with primary anti-EAAT2 antibody (Abcam ab#203130; 1.0 mg/ml at 1:1000 dilution) or primary anti-GAPDH (Abcam ab#9485; 1.0 mg/ml at 1:500 dilution). The blots were then incubated for 1 h with horseradish peroxidase-conjugated anti-rabbit IgG (1:40,000). Control western blot experiments to confirm antibody specificity were performed in the presence of secondary antibody only (i.e., absence of primary antibody). Membranes were developed using enhanced chemiluminescence (Super Signal West Pico, Thermo-Fisher). Bands were quantitated using ImageJ software (Wayne Rasband, Research Services Branch, National Institute of Mental Health, Bethesda, MD) and normalized to GAPDH. A commercially available cell lysate prepared from primary cultures of human astrocytes (ThermoFisher Scientific, Waltham, MA) was used as a positive control.

For western blot experiments on tissue specimens collected from frozen dorsal hippocampi, samples were homogenized in chilled T-PER lysis buffer (Cat. #: 78510, Thermo Fisher Scientific) supplemented with protease inhibitors (Cat. #: 04693116001, Roche, Basel, Switzerland) and phosphatase inhibitors (Cat. #: 04906837001, Roche) and centrifuged at 15,000 × g at 4°C for 15 min. The supernatants were collected, introduced to protein concentration assay using a BCA kit (Cat. #: 23225, Thermo Fisher Scientific) and adjusted to the same concentration. The supernatants (10 μg of total protein) were diluted in sample buffer (Cat# S3401, Sigma-Aldrich) with the omission of 2-mercaptoethanol and mildly heated at 65 °C for 10 min to avoid protein aggregation. The protein mixtures were resolved in polyacrylamide gels (8%) at 110 V for 2 h and transferred to PVDF membranes (Cat. #: IPVH00010, Merck-Millipore, Burlington, MA, USA). The membranes were blocked with 5% skim milk and hybridized with proper primary antibodies (SLC1A2: Cat. #: 3838, Cell Signaling Technology, Danvers, MA, USA; β-actin: Cat. #: A5441, Sigma-Aldrich) for 16 h at 4°C. After washing, the membranes were subsequently hybridized with proper horseradish peroxidase (HRP)-conjugated secondary antibodies (goat anti-mouse IgG: Cat. #: 115-035-166; goat anti-rabbit IgG: Cat. #: 111-035-144, Jackson ImmunoResearch, West Grove, PA, USA). The bound antibodies were detected using an enhanced chemiluminescence detection kit (Cat. #: WBKLS0500, Merck-Millipore) and X-ray film (Cat. #: Super RX, Fujifilm, Tokyo, Japan). Relative protein expression was estimated by normalizing with the β-actin level. The band densities were analyzed using ImageJ software (v2.0.0-rc-69/1.52p, U.S. National Institutes of Health, Bethesda, MD, USA). For re-probing, the membranes were incubated with a stripping buffer containing 2% SDS, 62.5 mM Tris, and 0.8% 2-mercaptoethanol for 20 min at 55 °C to remove the bound antibodies. The dilution ratios of antibodies and the image exposure time were tested to confirm that the luminescence signals were within the linear range of detection.

**12. [^3^H]Glutamic Acid Uptake Assay**

For cellular uptake experiments, cells were seeded into 24-well plates with a cell density of 2 x 10^4^ cells/cm^2^. All transport experiments were performed by using Hanks’ balanced salt solution (HBSS; transport buffer) containing 1.3 mM CaCl_2_, 0.49 mM MgCl_2_, 0.41 mM MgSO_4_, 5.3 mM KCl, 0.44 mM KH_2_PO_4_, 138 mM NaCl, 0.34 mM Na_2_HPO_4_, and 5.6 mM D-glucose. The buffer was supplemented with 25 mM HEPES, pH 7.4, and 0.01% bovine serum albumin to reduce the binding of radiolabeled compounds to assay plates. Confluent monolayers of HPMs were preincubated with transport buffer (pH 7.4) at 37°C for 30 min, and then incubated in transport buffer containing 1.0 μCi/ml [^3^H]glutamic acid (PerkinElmer Life Sciences, Boston, MA). To evaluate the effect of a SLC1A2 transport inhibitor (I.e, transporter specificity), 10 µM *N*-[4-(2-Bromo-4,5-difluorophenoxy)phenyl]-L-asparagine (WAY213613) was added both to the preincubation buffer and to the radioactive transport buffer. Stock solutions of WAY213613 (10 mM) were prepared in 50:50 HBSS:DMSO. In our transport experiments, the total concentration of DMSO in culture media did not exceed 0.1%. For inhibition experiments, cells were incubated with preincubation buffer containing WAY213613 for 10 min prior to adding transport buffer. At the desired time interval, the radioactive medium was removed and cells were washed twice with ice-cold NaCl (0.16 M) and solubilized in 1% Triton-X-100 at 37°C for 30 min. The content of each well was collected and mixed with 2.0 ml of Optiphase™ liquid scintillation cocktail (PerkinElmer) and the total radioactivity was measured with a model 1450 liquid scintillation counter (PerkinElmer). In each experiment, “zero time” uptake (i.e., background) correction was applied to correct for the nonspecific binding and variable quench time, estimated by the retention of radiolabeled compound in the cells after a minimum (zero) time of exposure, determined by removed the radiolabeled buffer immediately after its introduction into the well, followed by two washes with ice-cold HBSS and collection of cells for liquid scintillation counting. Uptake of the radiolabeled glutamic acid was normalized to total cellular protein content per well, which was measured by using Bradford’s method with BSA as the standard.

**13. Administration of Riluzole to 3xTg-AD mice**

**13.1) Animals**

All experiments were performed following the National Institutes of Health Guidelines for Animal Research (Guide for the Care and Use of Laboratory Animals) and approved by the National Cheng Kung University (NCKU) Institutional Animal Care and Use Committee (approval reference number: 106013). 3xTg-AD mice [B6;129-Tg(APPSwe, tauP301L)1Lfa Psen1tm1Mpm/Mmjax] exhibiting tau and amyloid pathologies were obtained from the Jackson Laboratory (Bar Harbor, ME, USA) and maintained in National Cheng Kung University Laboratory Animal Center (NCKULAC, Tainan, Taiwan) accredited by AAALAC. Since the AD pathologies are more stable and severe in females than in males[91], only female mice were used in this study. Mice were genotyped using protocols provided by The Jackson Laboratory. Mice were housed in ventilated cages (2-3 mice per cage) in a humidity- (55 ± 10%) and temperature-controlled (24 ± 1°C) specific-pathogen-free breeding unit of NCKULAC on a 12-h light/12-h dark cycle (light on at 7 AM). They had free access to water and diet. The housing environment and animal health were monitored by qualified staff and veterinaries. All experimental procedures were performed during the light cycle. This study included 20 15-month-old female 3xTg-AD mice which were randomly assigned to vehicle and riluzole groups (10 mice pre group) with computer-based randomization.

**13.2) Riluzole treatment**

Mice in the riluzole group were injected once daily with riluzole (RLZ; Cat. #: 1604337, Sigma-Aldrich, St. Louis, MO, USA; 4 mg/kg dissolved in normal saline containing 1% DMSO, i.p.) for 4 weeks. The mice received daily injections (i.p.) of an equal volume of saline containing 1% DMSO (Cat. #: D2650, Sigma-Aldrich) served as vehicle controls.

**13.3) Animal specimen preparations**

One day after termination of riluzole treatment, mice were anesthetized with Zoletil 50 (75 mg/kg, i.p.; Virbac, Carros, France) and transcardially perfused with chilled PBS (Cat. #: 10010023, Thermo Fisher Scientific, Waltham, MA, USA). Brains were quickly removed and divided into two hemispheres. The dorsal hippocampi were dissected from the left hemispheres, immediately immersed in liquid nitrogen, and stored at -80°C for further Western Blots and ELISA assays. The right hemispheres were post-fixed with 4% paraformaldehyde (Cat. #: 158127, Sigma-Aldrich) prepared in 0.1M phosphate buffer (Cat. #: P3619, Sigma-Aldrich) for 24 h and dehydrated daily with gradually increasing concentrations (10, 20, 30, and 35%, twice per concentration) of sucrose (Cat. #: S9378, Sigma-Aldrich) solutions prepared in 0.1 M phosphate buffer (Cat. #: P3619, Sigma-Aldrich). The dehydrated brains were embedded in the cutting compound (Cat. #: 3801480, Leica Biosystems, Wetzlar, Hessen, Germany) and prepared into 25-µm coronal sections using a cryostat (Model: CM1950, Leica Biosystems) for further immunohistochemistry.

**13.4) Aβ quantification**

The concentrations of soluble Aβ1-40 and Aβ1-42 in the dorsal hippocampi of mice were determined using commercial ELISA kits [human Aβ (1-40) ELISA kit: KHB3481; human Aβ (1-42) ELISA kit: KHB3441, Thermo Fisher Scientific Inc.] according to the manufacturer’s instructions.

**13.5) Immunohistochemistry**

Coronal brain sections (25 μm) containing dorsal hippocampus (from 1.46 mm to 2.18 mm posterior to the Bregma) were washed with TBS (Cat. #: R017R.0000, Thermo Fisher Scientific) containing 0.3% Triton X-100 (Cat. #: X100, Sigma-Aldrich) (TBS-T), incubated with TBS-T containing 3% H_2_O_2_ (Cat. #: H1009, Sigma-Aldrich) for 20 min at room temperature, blocked with 3% normal goat serum (Cat. #: S26-M, Sigma-Aldrich) prepared in TBS-T for 1 h at room temperature and probed with primary antibodies against human Aβ [clone 4G8: 1:200 dilution (Cat. #: SIG-39200, BioLegend, San Diego, CA, USA) + clone 6E10: 1:200 dilution (Cat. #: SIG-39300, BioLegend)], Tau (1:500 dilution, Cat. #: MAB3420, Merck-Millipore), phosphorylated Tau (clone AT8, 1:500 dilution, Cat. #: MN1020, Thermo Fisher Scientific), phosphorylated Tau^Thr181^ (1:500 dilution, Cat. #: 12885, Cell Signaling Technology), and phosphorylated Tau^S396^ (1:500 dilution, Cat. #: 9632, Cell Signaling Technology) for 16 h at room temperature. After washing to remove unbound antibodies, brain sections were incubated with HRP-conjugated secondary antibodies (1:500 dilution, goat anti-mouse IgG: Cat. #: 115-035-166; goat anti-rabbit IgG: Cat. #: 111-035-144, Jackson ImmunoResearch) for 2 h at room temperature, followed by TBS-T washes, and incubated with the chromogen, 3,3’-Diaminobenzidine (Cat. #: D12384, Sigma-Aldrich). After dehydration with ethanol solutions, sections were mounted with the xylene-based mounting medium (Cat. #: 3801730, Leica Biosystems).

**13.6) Image capture and analyses**

Immunohistochemical images were captured by an optical microscope (Model: Axio Imager A1, Carl Zeiss, Oberkochen, Germany) equipped with a digital camera (Model: Axiocam 305 Color, Carl Zeiss). The immunoreactive signals were identified by measuring the signal intensity higher than the background threshold using the ImageJ software (v2.0.0-rc-69/1.52p, U.S. National Institutes of Health, Bethesda). The background intensity threshold was fixed and applied to all sections. Five sections from each animal were determined, and the average was presented as a single data point.

**14. Statistical Analysis**

All numerical data are expressed as mean ± standard deviation. Statistical analyses and graph plotting were performed using the Prism software (v. 9.0a, GraphPad Software Inc., San Diego, CA, USA). Significance was set at *p* < 0.05. The D’Agostino-Pearson normality test was adopted to estimate the assumption of normality. For datasets with normal distribution, the unpaired, two-tailed Student’s t-test was used to compare means between two groups. For non-normal distributed datasets, the Mann-Whitney U test was used.
